# In vivo and in silico alpha-synuclein propagation dynamics: The role of genotype, epicentre, and connectivity

**DOI:** 10.1101/2025.07.18.665565

**Authors:** Stephanie Tullo, Janice SH Park, Shady Rahayel, Daniel Gallino, Megan Park, Kristie Mar, Ying-Qiu Zheng, Esther del Cid-Pellitero, Edward A. Fon, Wen Luo, Irina Shlaifer, Thomas M. Durcan, Bratislav Misic, Alain Dagher, Gabriel A. Devenyi, M. Mallar Chakravarty

**Affiliations:** Integrated Program in Neuroscience, McGill University, Montreal, Quebec, Canada; Cerebral Imaging Center, Douglas Mental Health University Institute, McGill University, Verdun, Quebec, Canada; Department of Medicine, University of Montreal, Montreal, Quebec, Canada; Center for Advanced Research in Sleep Medicine, Hôpital du Sacré-Coeur de Montréal, Montreal, Quebec, Canada; Wellcome Centre for Integrative Neuroimaging, University of Oxford, Oxford, United Kingdom; Department of Neurology and Neurosurgery, Montreal Neurological Institute, McGill University, Montreal, Quebec, Canada; McGill Parkinson Program, Neurodegenerative Diseases Group, Department of Neurology and Neurosurgery, Montreal Neurological Institute-Hospital, McGill University, Montreal, Québec, Canada; Early Drug Discovery Unit, Montreal Neurological Institute, McGill University, Montreal, Quebec, Canada; Department of Biological & Biomedical Engineering, McGill University, Montreal, Quebec, Canada; Department of Psychiatry, McGill University, Montreal, Quebec, Canada

## Abstract

Neurodegeneration observed in synucleinopathies, like Parkinson’s disease (PD), are hypothesized to be a consequence of the progressive accumulation and spread of misfolded alpha-synuclein (aSyn) throughout the brain. Here, we study the generalizability of this hypothesis across multiple biologically-relevant factors including genotype, aSyn species, and regional vulnerability (i.e.: does the introduction of aSyn into different brain regions have a similar impact) using a detailed longitudinal brain (using magnetic resonance imaging; [MRI]) and behavioural approaches. We first examined wild-type and M83 A53T hemizygous transgenic (engineered to over-express human aSyn) C57BL/6 x C3H mice receiving striatal inoculation with human or mouse preformed fibrils (PFFs) (n=89 mice at the last time point; n=687 MRI). Longitudinal analyses demonstrated a time-dependent increase in network-like atrophy and motor deficits, generalized across genotype and PFF species. We further derived latent dimensions relating brain-behaviour relationships revealing a pattern of sex- and genotype-relevant atrophy. Overall, atrophy was most prominent in M83 mice with mouse PFFs, while human PFFs or wild-type hosts showed attenuated effects. Changing the inoculation site to the hippocampus, a major connectivity hub, revealed differential regional vulnerability in the form of localized atrophy. Computational models previously validated in clinical PD further indicated regional vulnerability with increased predictive ability for atrophy patterns associated with striatal compared to hippocampal inoculation. Our findings reveal atrophy resulting from aSyn spreading is generalizable across genotype and PFF species, but not disease epicentres, emphasizing the role of regional vulnerability in disease progression.

## 1. Introduction

The aggregation and misfolding of alpha-synuclein (aSyn) protein (Goedert et al. 2013; Braak et al. 2003) is considered central to the pathology of synucleinopathies like Parkinson’s disease (PD). While the mechanisms underlying the aSyn pathological cascade remain elusive, evidence suggests a pivotal role for the prion-like spreading of pathological aSyn in disease progression and downstream atrophy (Bétemps et al. 2014; Desplats et al. 2009; Fares et al. 2016; Froula et al. 2019; Kordower et al. 2008; Li et al. 2008; Luk et al. 2012a; Luk et al. 2012b; Masuda-Suzukake et al. 2013, 2014; Mougenot et al. 2012; Panattoni et al. 2018; Sacino et al. 2014; Tapias et al. 2017; Volpicelli-Daley et al. 2014; Watts et al. 2013). This process involves the uptake of pathological aSyn by healthy neurons, triggering the misfolding and aggregation of endogenous aSyn, and subsequent spread throughout the entire brain (Aguzzi and Rajendran 2009; Clavaguera et al. 2009; de Calignon et al. 2012; Desplats et al. 2009; Hansen et al. 2011; Meyer-Luehmann et al. 2006; Prusiner 2012; Warren et al. 2013). However, the extent to which the progressive spreading of pathogenic aSyn is related to brain-region specific atrophy that underlies symptomatology is still unknown. Understanding the mechanisms underlying aSyn propagation and its effects on brain circuits, as well as its downstream consequences on behaviour and disease worsening, is critical for elucidating synucleinopathy-associated pathogenesis. This understanding can, in turn, be used to understand the efficacy of therapeutic strategies.

To instigate disease propagation from a known epicentre, recent studies have seeded aSyn preformed fibrils (PFF) into brain regions of both wild-type mice of various strains and transgenic mice overexpressing human A53T aSyn (M83 mice) (Giasson et al. 2002). These models recapitulate key features of synucleinopathies, including progressive motor impairments and neuroanatomical pathology, including atrophy (Froula et al. 2019; Luk et al. 2012a; Luk et al. 2012b; Masuda-Suzukake et al. 2013, 2014; Mougenot et al. 2012; Sacino et al. 2014; Tullo et al. 2023, 2025; Watts et al. 2013). We recently demonstrated that this model recapitulates key neuroanatomical and behavioural features (Tullo et al. 2023) relevant to human synucleinopathies (Zeighami et al. 2015) Additionally, more recently, we demonstrated a differential resilience to disease progression between the sexes for M83 mice injected with human PFF, with a faster progression in males (Tullo et al. 2025). Based on these findings, it is clear that the generalizability of the aSyn spreading hypothesis requires further investigation across other biological factors that may impact disease progression, namely genotype (genetic predisposition), PFF species, and specific regional vulnerability (i.e.: disease epicentre). Examining these factors allows for a detailed examination of the prion-like transmission hypothesis.

This hypothesis has been used to model and predict aSyn spreading and aSyn-mediated neurodegeneration in both animal models and clinical PD. These models operate on the assumption that both axonal connectivity and the spatial distribution of the expression of genes (Hawrylycz et al. 2012; Lein et al. 2007; Ng et al. 2009; Oh et al. 2014; Kuan et al. 2015) underlying aSyn production and clearance (Zheng et al. 2019; Rahayel et al. 2022b; Rahayel et al. 2023; Rahayel et al. 2022a) act as constraints on both aSyn propagation and downstream neurodegenerative processes. This model was recently used to predict misfolded aSyn burden in a mouse model across multiple seed regions (Rahayel et al. 2022a). However, the relationship between aSyn propagation and atrophy remains unclear, and the mechanistic dependency on disease epicentre, and consequently regional vulnerability, could be further tested using these sophisticated computational methods.

Here, we investigated the generalizability of the prion-like aSyn spreading hypothesis by expanding on previous work from our group (Tullo et al. 2023, 2025). In our previous work we longitudinally examined brain and behavioural phenotypes in the M83 mouse model inoculated with human PFFs in the dorsal striatum. Here, we expand on this approach as a means to gain mechanistic insight on how PFF species (mouse or human PFF) and mouse genotype (M83 hemizygous versus WT) modulate aSyn spreading dynamics leading to downstream neurodegeneration captured by magnetic resonance imaging (MRI). We further examine the role of regional vulnerability, as indexed by differential atrophy patterns, by varying the injection site (striatum versus hippocampus, another highly connected region of the brain). Finally, we employed the computational model mentioned above to assess if seed point specific neurodegenerative patterns could be predicted using knowledge regarding the mouse brain structural connectome and the transcriptomic profile related to aSyn production across both disease epicentres. While our findings affirm the nigro-striatal pathway’s vulnerability across PFF species and mouse strains, our hippocampal experiments revealed striking regional disparities across both *in vivo* and *in silico* experiments. These findings of differential regional vulnerability may provide critical insights into the mechanisms driving synucleinopathy-induced neurodegeneration and pave the way for the development of effective therapeutic interventions using preclinical and computational models.

## 2. Results

To directly test the robustness of the aSyn prion-like spreading hypothesis and its relationship to brain atrophy and motor impairments, we modeled synucleinopathy-induced neurodegeneration over the disease time course in mice. Adult (11-12 weeks old) hemizygous M83 aSynA53T transgenic mice and wild-type (WT) littermates (C57BL6/C3H) were injected in the right caudoputamen via intracerebral injection with either aSyn preformed fibrils (PFF) or phosphate buffered saline (PBS) (n∼8 mice per genotype/injection group/sex at the last time point; subject numbers at each time point for each condition in Supplementary Table 1). Here, we used a similar longitudinal approach to examine neuroanatomical changes, while assessing symptomatology and disease progression within the same mice (as previously shown in other models of disease: (Borg and Chereul 2008; Gallino et al. 2019; Guma et al. 2021, 2020; Johnson et al. 2007; Kong et al. 2018; Pagani et al. 2016; Rollins et al. 2019; Tullo et al. 2025)), to determine the influence of host genotype and PFF species on spreading patterns, disease progression, and survival. We characterized differences in MRI-derived neurodegeneration, motor impairments, and survival rates in WT and M83 hemizygous mice that received either PBS, human [Hu-] or mouse [Ms-] PFF inoculation (Volpicelli-Daley et al. 2014); characterization: Supplementary Figure 1) across disease progression (−7, 30, 90 and 120 days-post-injection; dpi). An overview of the experimental design and timeline is provided in Figure 1.

**Figure 1.**
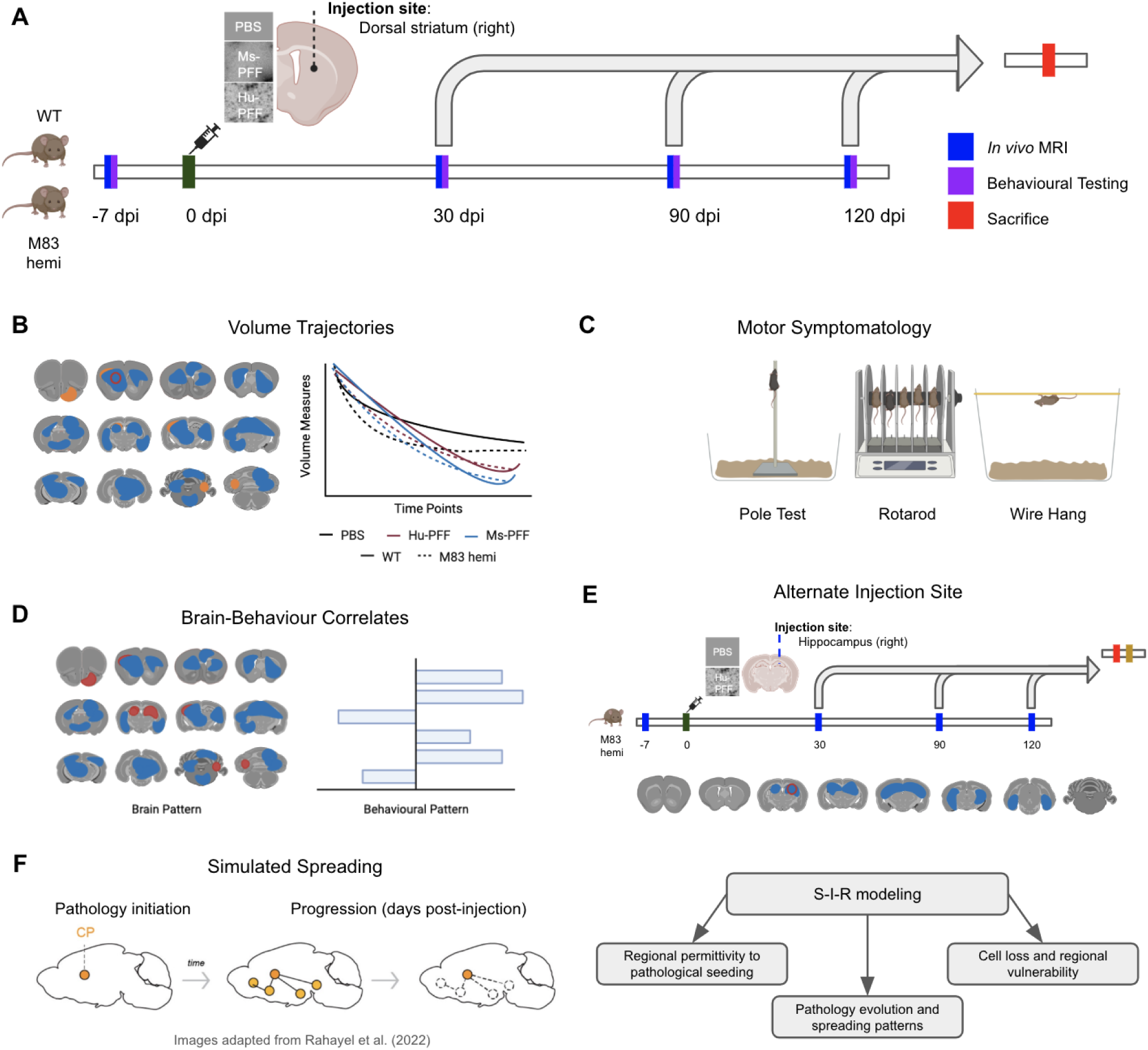
Overview of workflow. [A] Experimental timeline of mouse experiments where wild-type (WT) and M83 hemizygous mice were injected with either phosphate buffered saline (PBS), human (Hu-) or mouse (Ms-) preformed fibrils (PFF) in the right striatum. Imaging and behavioural testing was administered at four time points: −7, 30, 90 and 120 dpi. Mice were siphoned off at each post-injection time point. [B] Neuroanatomical changes over the course of the experiment were assessed to examine MRI-derived PFF-induced atrophy. [C] Motor symptomatology was assessed based on the performance of the mice on the pole test, rotarod and the wire hang test at each of the four time points. [D] Correlative relationship between MRI-derived brain pathology and demographic measures (such as motor symptomatology, injection group, sex, genotype) was examined using partial least squares analysis (PLS). [E] Experimental timeline of the second mouse experiment, such that M83 hemizygous mice received an injection of PBS or Hu-PFF in the right dentate gyrus of the hippocampus to assess aSyn spreading via an alternate disease epicenter. Neuroanatomical changes over time were similarly examined here. [F] Computational modelling of aSyn propagation was generated using the S-I-R model adapted from Rahayel et al. (2022a; 2022b) and Zheng et al. (2019). Accuracy of the *in silico* model was assessed via comparison of simulated atrophy to empirical atrophy derived from the mouse experiments for both the caudoputamen and hippocampal injection site.

### 2.1. Disease progression and overt motor impairment is present across genotype and PFF species

Striatal PFF inoculation resulted in time-dependent motor impairments and disease progression that was further modulated by genotype and PFF species. We observed dramatic reductions in survival of M83 PFF-injected mice, regardless of PFF species, compared to all other groups by 120 dpi with significantly lower rates of survival for the M83 Ms-PFF compared to the M83 Hu-PFF-injected group (p=0.00271; HR = 0.305; CI = 0.14 - 0.66; ∼70% higher hazard compared to M83 Hu-PFF [Figure 2B]). No injection group survival sex differences were observed in M83 Ms-PFF-injected mice (Figure 2C) (females: n=13/23; males: n=10/23). However, for the M83 Hu-PFF-injected mice, only male mice reached a humane endpoint prior to the experimental endpoint (n=9/23); no female mice reached a humane endpoint during this experiment timeframe (n=0/19).

**Figure 2.**
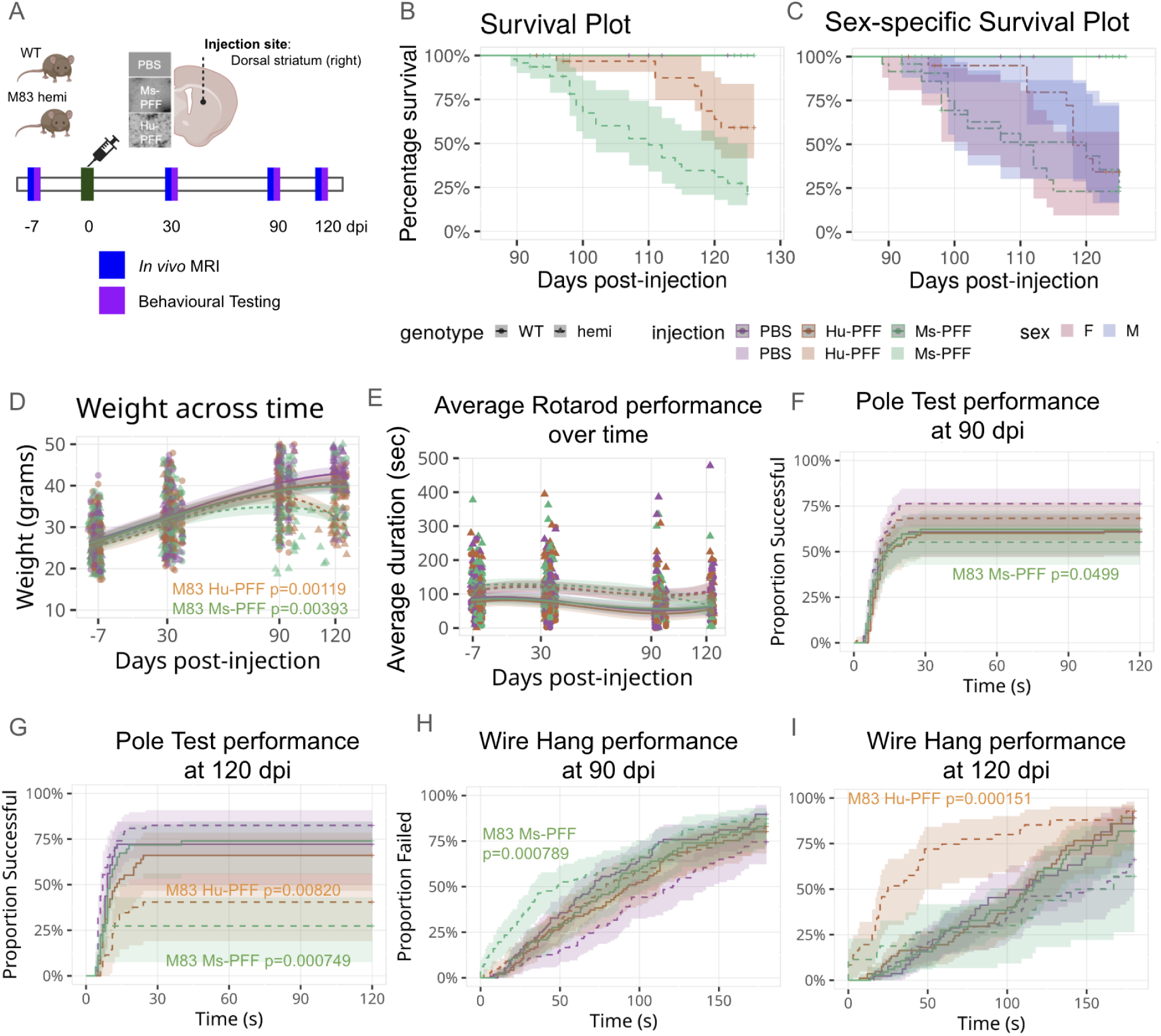
Disease progression and motor symptomatology. [A] Experimental timeline for WT and M83 hemizygous mice injected with either PBS, Ms-PFF, or Hu-PFF in the right striatum, with MRI and behavioural testing conducted at −7, 30, 90, and 120 dpi. [B] Percent survival plotted across dpi, with the experimental endpoint of ∼130 dpi. M83 PFF-injected mice showed reduced survival, particularly for the M83 Ms-PFF-injected mice (p=5.50e-5). [C] Sex differences in survival rates for M83 Hu-PFF-injected mice, with lower rates in males (p=0.000411). No significant sex differences in survival rates within M83 Ms-PFF injected mice. [D] Weight trend across disease progression showed significant weight loss for M83 Ms-PFF and Hu-PFF-injected mice, with weight loss beginning around 90 dpi (p=0.00119; p=0.00393 respectively) (compared to WT PBS-injected mice). [E] Average rotarod performance across time showed no significant differences between injection groups and genotype across the time points. [F,G] Pole Test performance at 90 dpi [F] and 120 dpi [G]. Pole Test performance at 90 and 120 dpi, indicating worse performance for M83 Ms-PFF mice at both time points (p=0.0499; p=0.000749), and for M83 Hu-PFF at 120 dpi (p=0.00820). [H,I] Wire hang test performance at 90 dpi [H] and 120 dpi [I], performing worse for M83 Ms-PFF at 90 dpi (p=0.000789) and M83 Hu-PFF at both time points (p=0.00691; p=0.000151) compared to WT PBS mice. Purple colour denotes PBS-injected mice, orange colour denotes Hu-PFF-injected mice, and green colour denotes Ms-PFF-injected mice. Line type and shape of points were used to denote each of the genotypes: solid line and round point for wild-type, and dashed line and triangular points for M83 hemizygous mice.

Upon starting blinded daily observation of mice ambulation in their home cages at 80 dpi, we observed motor impairments (including freezing gait and impairments in rearing behaviours) more frequently for M83 (Hu: n=36/42; Ms: n=45/46) versus WT mice (Hu: n=21/36; Ms: n=21/34; Fisher’s Exact Test for Count Data: p= 2.87e-6; Supplementary Figure 2A-B). Pairwise comparisons between each of the four groups were performed, and while we observed a significant difference between M83 Ms-PFF and M83 Hu-PFF (p=0.0507), this significance did not survive multiple comparisons correction (Supplementary Figure 2B). Other significant pairwise comparisons after FDR correction are available in Supplementary Figure 2B. Sex differences in motor symptomatology are detailed in Supplementary Figure 2.

Quantitative assessment of disease progression via weight tracking and motor performance reveal a similar trend of most severe impact for M83 PFF-injected mice, regardless of PFF species. We observed a significant injection group by genotype by time interaction on weight across four time points (−7, 30, 90 and 120 dpi) with an inverted U-shaped cubic trajectory for the M83 Hu-PFF-injected mice (df=952.274; p=0.00119) and a linear decline for M83 Ms-PFF mice (df=980.690; p=0.00393) compared to WT PBS-injected mice (Figure 2D). Furthermore, M83 Ms-PFF demonstrated a faster rate of weight loss versus M83 Hu-PFF mice (df=957; p=0.0001). Given the known differences in weight between male and female mice, this relationship was further examined in a sex-specific manner (Supplementary Figure 2F).

In sum, M83 PFF-injected mice demonstrated disease progression that led to weight loss and motor impairment progression leading to a humane endpoint. This demonstrates that, as expected, the M83 genotype accelerates disease progression brought on by PFF inoculation.

### 2.2. PFF species independence on worsening motor performance for M83 hemizygous mice

Rotarod, pole test and wire hang were similarly assessed at the four time points. We observed no differences in average rotarod performance across injection groups or genotypes over time (Figure 2E). We examined both failure/success rate as well as performance duration for each task cross-sectionally at each timepoint for the pole test and wire hang test. Consistent with the above observations of motor impairments, we observed significant worsening of performance for M83 PFF-injected mice at the 90 dpi indexed by slower descents and lower rates of successful completion (failures include exceeding a 120 second time limit or failing to grip and falling down the pole; p=0.0499; Hazard ratio [HR]=0.463; Figure 2F) as well as shorter hanging durations and higher rates of failure on the wire hang test (less than 10 seconds hanging on the third consecutive try; p=0.0.000789; HR=2.087; Figure 2G). At 120 dpi, only the M83 Hu-PFF injected mice performed significantly worse than WT PBS mice on the wire hang test (p=0.000151; HR=3.201; Figure 2H), while both Ms- and Hu-PFF injected M83 mice performed significantly worse on the pole test (p=0.00820 for M83 Hu-PFF mice and p=0.000749 for M83 Ms-PFF mice; Figure 2I). No sex differences were observed for any of these measures (Supplementary Figure 3).

While striatal PFF inoculation resulted in time-dependent motor impairments and disease progression with differing rates based on the genotype of the mice and species of PFF injection, nonetheless, the phenotype is robust to genotype and species inoculum.

### 2.4. Widespread neuroanatomical changes over time for striatal PFF-injected mice

Striatal PFF inoculation induces widespread neuroanatomical changes over time regardless of mouse genotype. Voxel-wise volume measures were obtained from the Jacobian determinants derived from the deformation fields of the nonlinear transformations (Chung et al. 2001) generated from longitudinally optimized group-wise registrations (deformation-based morphometry; DBM; (Germann et al. 2025)) of T1-weighted MRI (Bruker 7T MRI BioSpec 70/30 USR with 1H Quadrature transmit/receive MRI CryoProbe; 100 μm^3^ isotropic voxels; equivalent to the standard 1 mm^3^ voxels in human MRI) at −7, 30, 90, and 120 dpi for each subject. We observed significant changes in brain anatomy (steeper rates of change; assessed using linear mixed effects models examining genotype by injection group over time) in PFF mice, regardless of PFF species (when comparing to WT PBS mice, as well as within the M83 mice groups) at a strict FDR threshold of 1% (Figure 3A-E). In line with our previous finding (Tullo et al. 2025), the greatest statistical differences were observed in the areas with known connections to and from the injection site (right striatum) (Figure 3H), such as the right primary motor area (Figure 3I), the right substantia nigra pars compacta (Figure 3G), and the periaqueductal gray (Figure 3F). Notably, the steepest rate of volumetric decline was observed for the M83 Ms-PFF injected mice (Figure 3B, E). When examining the deformation map for M83 Hu-PFF injected mice, similar areas showing volumetric decline are observed, however to a lesser degree and in fewer areas (Figure 3A, D). Voxels with steeper volume decline in M83 Ms-PFF-injected mice compared to M83 Hu-PFF-injected mice are explicitly shown in Figure 3C. Additional analyses examining sex differences, within injection genotype differences (M83 vs WT) and within genotype injection differences (PBS vs Hu vs Ms) are displayed in Supplementary Figures 4-9. These findings underscore the differential impact of aSyn spreading by highlighting the relevance of PFF species. Nonetheless, we observe a robust neurodegenerative phenotype across synucleinopathy models and PFF species.

**Figure 3.**
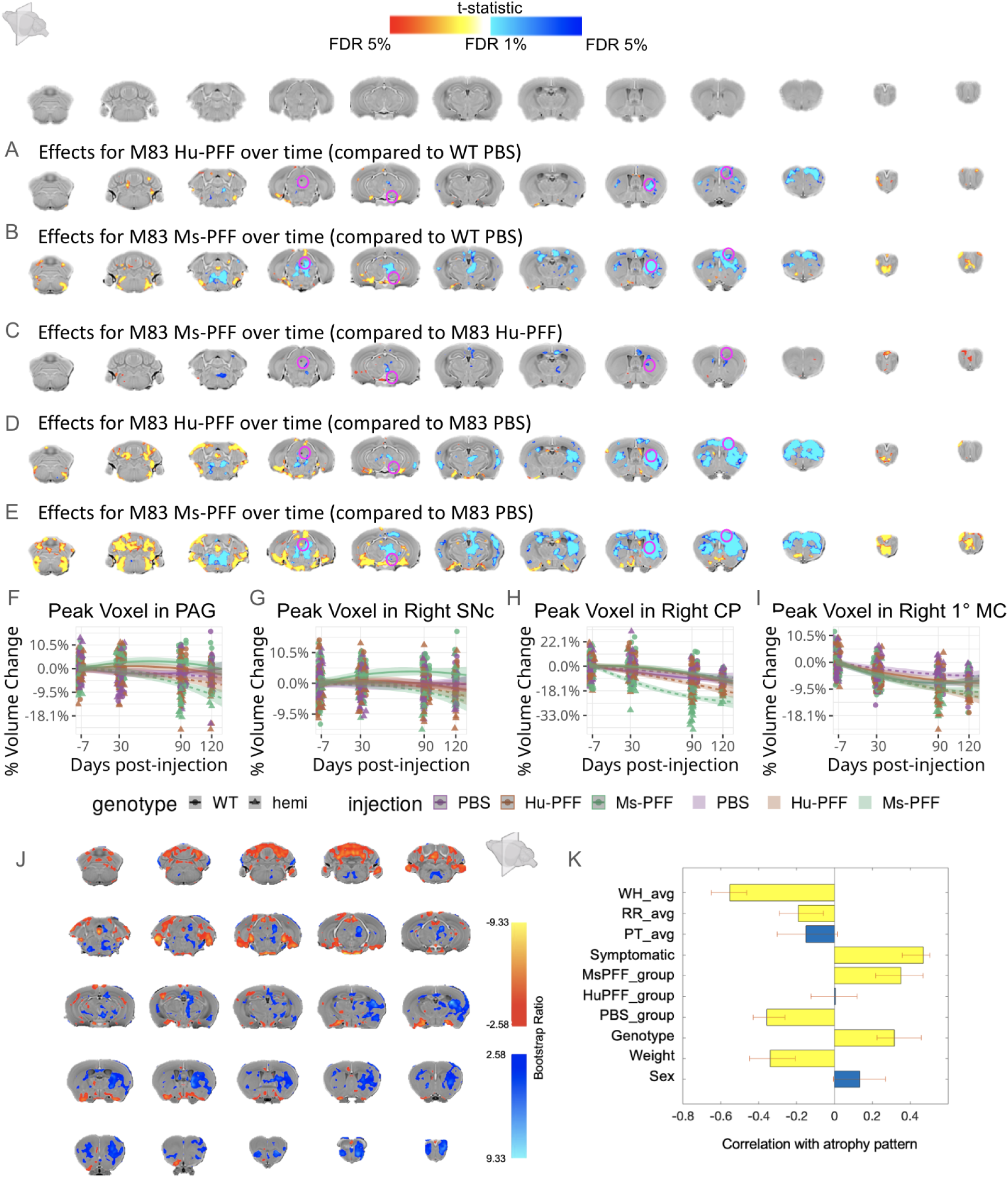
Brain-behavioural correlates of PFF-induced pathology and disease worsening. [A-E] Coronal slices of the mouse brain (from posterior to anterior) with t-statistical map overlay; demonstrating the effects of [A] M83 Hu-PFF injected mice or [B] M83 Ms-PFF-injected mice (compared to WT PBS mice), [C] M83 Ms-PFF compared to M83 Hu-PFF injected mice, [D] M83 Hu-PFF-injected mice and [E] M83 Ms-PFF-injected mice compared to M83 PBS mice. Colour map denotes significance level of voxel-wise volumetric trajectories over time with cooler colours denoting significant volumetric declines over time while warmer colours denote the opposite for the group of interest relative to the control group. Pink circles denote the peak voxel being plotted. [F-I] Plots of relative volume change (mm^3^) (compared to the mean volume per group of interest at the pre-injection time point; −7 dpi) over the four time points for a peak voxel in [F] the periaqueductal gray, [G] the right substantia nigra pars compacta (SNc), [H] the injection site (right caudoputamen (CP)), and [I] the right primary motor area (1 MC). Purple colour denotes PBS-injected mice, orange colour denotes Hu-PFF-injected mice, and green colour denotes Ms-PFF-injected mice. Line type and shape of points were used to denote each of the genotypes: solid line and round point for wild-type, and dashed line and triangular points for hemizygous mice. [J,K] Partial least squares (PLS) analysis results for the first latent variable (LV1). [J] Brain loading bootstrap ratios for the LV1 deformation pattern overlaid on the population average, with negative bootstrap ratios in orange-yellow (indicative of negative correlations with behavioural loadings [K]), and positive in blue (indicative of positive correlations with behavioural loadings [K]). Coloured voxels make significant contributions to LV1. [K] Behaviour weights for each behavioural measure included in the analysis showing how much they contribute and the direction of their correlation to the pattern of LV1. Singular value decomposition estimates the size of the bars whereas confidence intervals are estimated by bootstrapping. Bars with error bars that cross the 0 line (blue) are not considered (non-significantly associated). Abbreviations: WH_avg, average wire hang performance; RR_avg, average rotarod performance; PT_avg, average pole test performance.

### 2.5. Brain-behavioural latent dimensions of synucleinopathy-associated disease pathogenesis

To evaluate the impact of the spreading of misfolded aSyn concurrently using MRI and behavioural testing, partial least squares (PLS) analysis (Zeighami et al. 2019; McIntosh and Mišić 2013) was implemented to relate MRI-derived atrophy (voxel-wise volume difference between the −7 & 90 dpi peri-motor symptom onset time point) to symptom profile (sex, weight, genotype, injection, symptom score & motor performance) (n>=15 mice per genotype per injection group per sex), as done in previous work by our group (Guma et al. 2021). We observed a widespread pattern of brain pathology that was strongly correlated with the occurrence of overt motor symptomatology, lower weight, performing worse on the wire hang and rotarod test, as well as being M83 and having received a Ms-PFF injection (Figure 3J-K). This latent variable explained ∼34% of the covariance and was highly significant (p<0.0001; evaluated using permutation test; contribution of variables assessed with bootstrap resampling; (Zeighami et al. 2019; Patel et al. 2020)). Here we highlight a joint brain-behavioural latent dimension describing widespread patterns of volumetric decline in areas that are correlated to lower survival rates, shorter motor symptom duration, and overall poorer performance on the motor tasks for M83 Ms-PFF mice. This latent dimension describes inter-individual variation in brain atrophy that optimally maps onto a specific behavioural profile, genotype, and injection groups. The regions displaying MRI-derived atrophy are known to be highly connected to the striatum, suggesting that PFF-induced atrophy appears to be mediated by preferential aSyn propagation via structural connectivity, and in turn underlies this behavioural dimension.

### 2.6. Regional vulnerability is conferred by SNCA expression and not structural connectivity

After establishing that PFF inoculation in hemizygous mice results in an atrophy pattern resembling what has previously been observed in clinical groups (Tullo et al. 2023, 2025; Zeighami et al. 2015; Zheng et al. 2019), we sought to examine if atrophy patterns were closely linked to the structural connectome or the expression of genes that encodes aSyn (namely the SNCA gene). We first investigated the relationship between MRI-derived atrophy and structural connectivity measures provided by the Allen Institute’s Mouse Brain Connectivity Atlas (AMBCA). Viral tracing data from AMBCA (Oh et al. 2014), describing anterograde neuronal single synapse connectivity from 212 source regions (from right hemisphere), quantified connectivity strength measures between each of the source injection sites and its receiving target regions for 426 bilateral regions (Oh et al. 2014). Surprisingly, we obtained a relatively low correlation of r=0.017 (Figure 4A), suggesting that the atrophy cannot be predicted by structural connectivity alone. We next evaluated the correlation of MRI-derived atrophy with regional SNCA gene expression (Lein et al. 2007) with the prediction that regions exhibiting atrophy may have higher synucleinopathy-induced pathology as there are more proteins to convert to its pathogenic misfolded state. Notably, the correlation between such atrophy and *SNCA* expression was higher, with r=0.482 (Figure 4B). However, more likely, the combination of both structural connectivity and regional *SNCA* expression may have better predictive power of aSyn-induced neurodegeneration.

**Figure 4.**
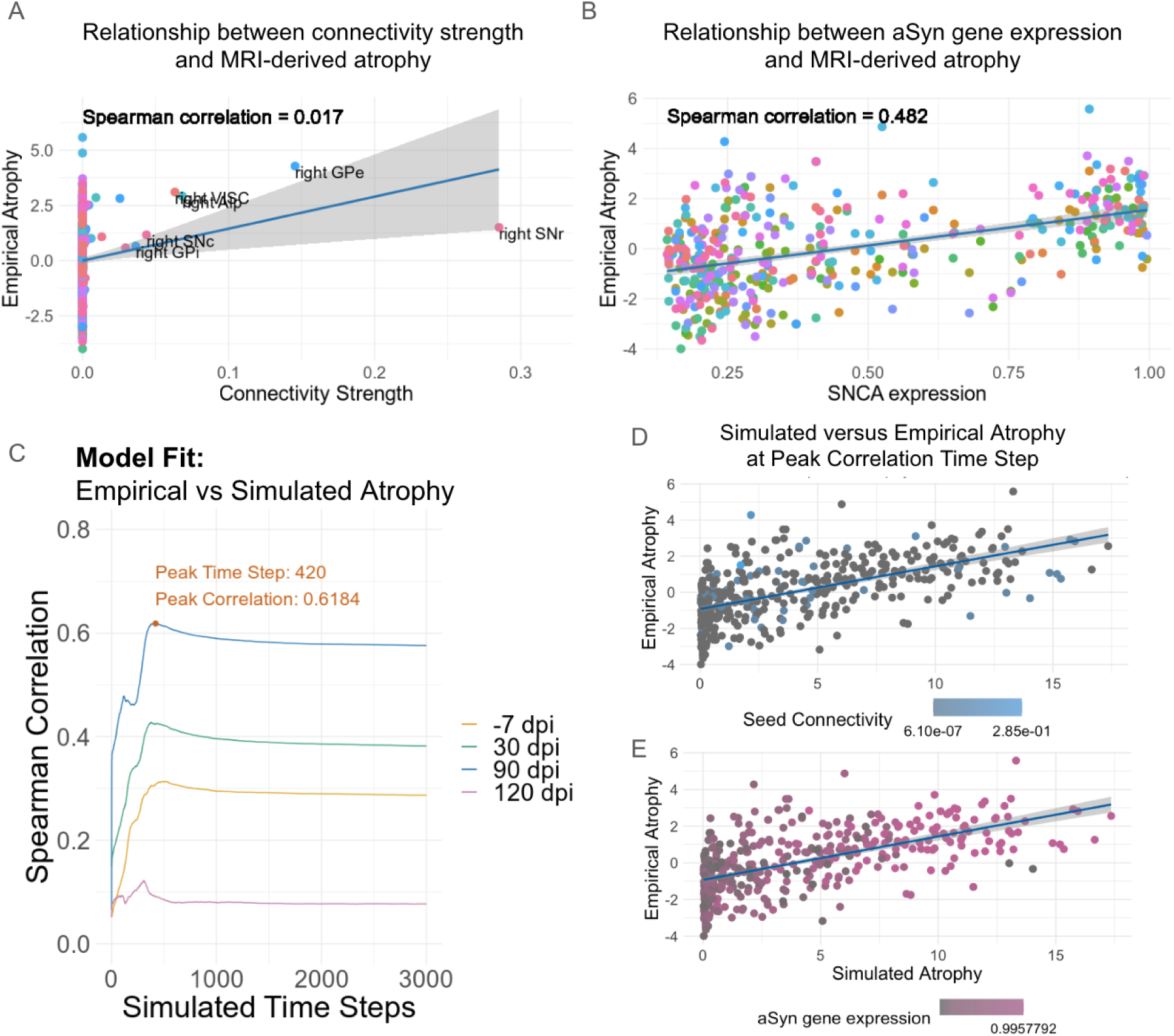
Simulated spread of aSyn propagation from striatal inoculation, utilizing both aSyn gene expression and structural connectivity, can accurately model empirical atrophy in a mouse model of synucleinopathy. [A] Spearman correlation between empirical atrophy and connectivity strength (r=0.017) was lower than [B] the correlation between atrophy and SNCA expression (r=0.482). Top correlated atrophy-connectivity regions include the right substantia nigra pars reticulata (SNr) and pars compacta (SNc), the right globus pallidus external (GPe) and internal (GPi) and the right Visceral area (VISC) and the right Agranular insular area posterior part (Alp). [C] Spearman correlations between simulated and empirical atrophy derived from M83 Ms-PFF (the most affected group) compared to WT PBS-injected mice at −7 (orange), 30 (green), 90 (blue) and 120 (pink) dpi. Correlations are shown as a function of simulation time. After reaching the peak value (r = 0.618), the model fit slightly drops and stabilizes. [D,E] Plots of regional pathology (empirical atrophy values at 90 dpi) versus simulated atrophy at the peak model fit (peak correlation time step). Points represent each region of the brain and is colour coded by either [D] its connectivity strength (blue gradient) with respect to the injection site seed, or [E] denotes its local aSyn gene (SNCA) expression (pink gradient) for each region of the AMBA.

### 2.7. Regional vulnerability simulated by computational model accurately predicts spread of atrophy from striatal seed

Simulated spread of aSyn propagation can accurately model empirical atrophy from striatal inoculation. Employing a computational model proven to accurately simulate atrophy in humans with synucleinopathies (Abdelgawad et al. 2022; Rahayel et al. 2022b; Zheng et al. 2019) provides a translational framework for elucidating the spatiotemporal progression of aSyn pathology and identifying factors which may mediate aSyn spread across the brain. This *in silico* modeling technique utilizing both *SNCA* expression and structural connectivity holds promise for understanding and predicting the complexity of aSyn propagation, recapitulating the spatiotemporal distribution of PFF-induced brain atrophy observed *in vivo*.

We simulated aSyn propagation and its associated neurodegeneration using a Susceptible-Infected-Removed (S-I-R) agent-based model, where aSyn agents can be in one of three states: “Susceptible” (normal), “Infected” (abnormal misfolded form), or “Removed” (metabolized protein). Synthesis rates of “Susceptible” agents were defined as proportional to *SNCA* expression, and agent propagation was based on the AMBCA. Interactions between “Infected” and “Susceptible” interacting agents transformed the latter into “Infected”. Simulated neurodegeneration was defined as one, regional atrophy (proportional to the accumulation of “Infected” agents) and two, deafferentation (induced by “atrophied” connected regions). Model-based simulated atrophy was compared to empirical atrophy using a Student’s t-test of the average Jacobian values normalized by region (n=426 regions) in the Allen Mouse Brain Atlas between the M83 Ms-PFF mice compared to WT PBS mice (where most severe disease phenotype was observed). We used Spearman correlations between simulated atrophy at every time step of the model with empirical atrophy values obtained at each of the four MRI time points (Figure 4E).

Comparisons between simulated and empirical atrophy revealed a peak correlation of r=0.618 at the 420th time step of the simulation and atrophy data at 90 dpi (Figure 4C). Fittingly, peak correlations were lower for earlier time points (−7 dpi: r=0.313; 30 dpi: r=0.428); however, even lower at 120 dpi, likely due to lower sample size as a result of lower survival rates for the M83 Ms-PFF mice (n=25 mice at 120 dpi vs n=83 mice at 90 dpi; r=0.122). At the peak correlation, simulated and empirical atrophy were examined with respect to connectivity strength from the injection site seed (Figure 4D) or *SNCA* expression (Figure 4E) for each region of the AMBCA. Despite capturing empirical atrophy patterns emanating from the caudoputamen using regional vulnerability measures such as gene expression and connectivity profile, whether these same principles generalize to non-striatal epicenters remains to be determined.

### 2.8. Brain wide pattern of pathology differs with hippocampal inoculation site

To further investigate the generalizability of aSyn spreading, we examined disease progression for hippocampal-injected mice. The rationale for choosing the hippocampus rested on its position as a highly connected network hub in the brain, that is not classically associated with PD brain atrophy. Further, this epicentre has been previously studied in previous studies and there is evidence in aSyn propagation resulting from PFFs injected into the hippocampus (Bétemps et al. 2014; Dues et al. 2023; Luk et al. 2012b; Rahayel et al. 2022a). However, the relationship between this propagation and downstream atrophy has yet to be examined. M83 hemizygous mice were injected with 2.5 μL Hu-PFF (or PBS) into the right dentate gyrus of the hippocampus (n∼8 injection group/sex at each time point). Similarly to the experimental design of the striatal injected mice, mice underwent *in vivo* MRI (T1-weighted images; 100 μm³ isotropic voxels) at −7, 30, 90 and 120 dpi (Figure 5A). Neuroanatomical measures, obtained using DBM, at the voxel level, were assessed to investigate whole brain volumetric change longitudinally using linear mixed effect models, as done previously for the striatum injection.

**Figure 5.**
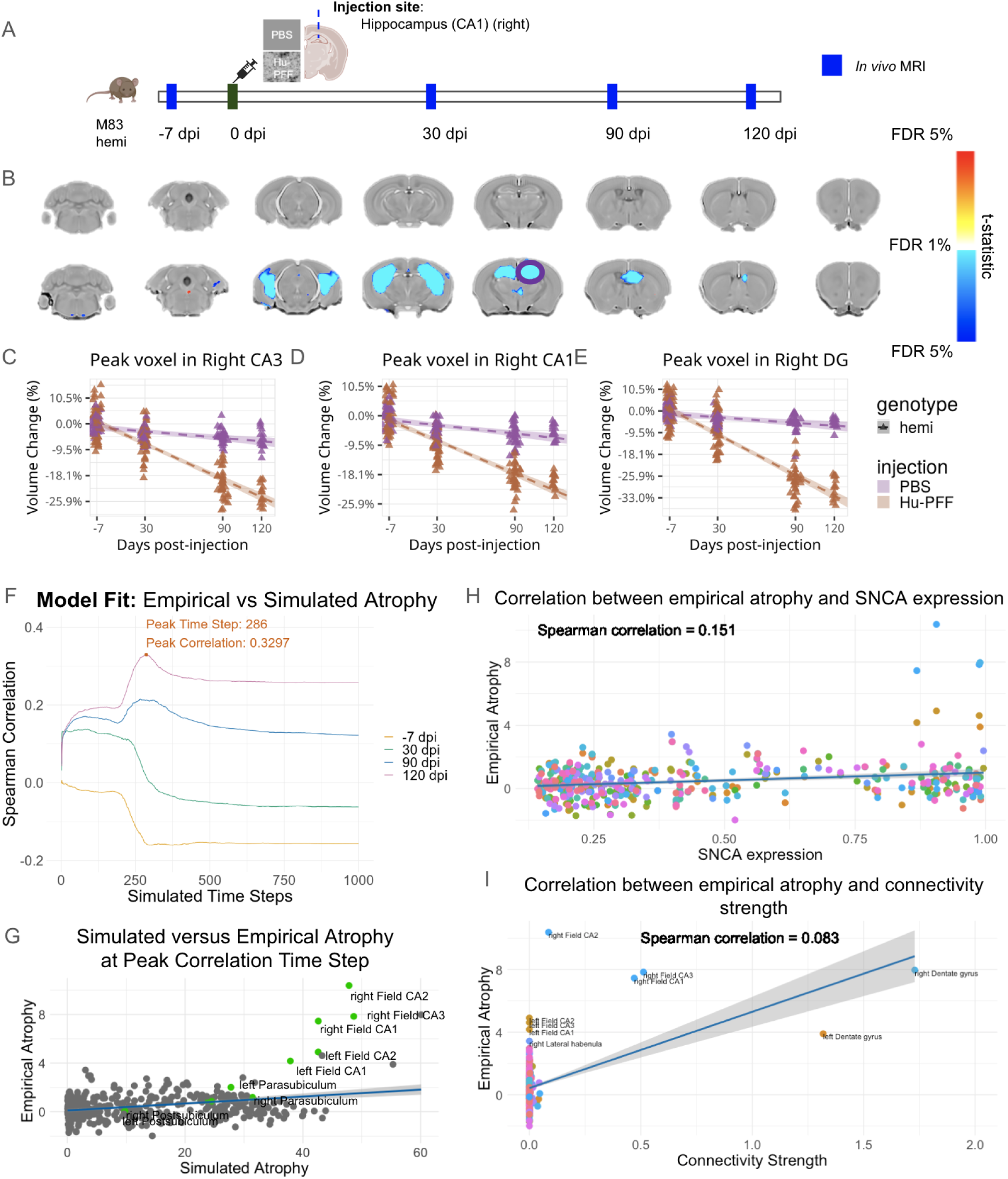
Hippocampal seeding of aSyn fibrils results in hippocampal-focused atrophy not well predicted by SNCA local gene expression and structural connectivity. [A] *In vivo* T1-weighted MRI (100 μm^3^ isotropic voxels; Bruker 7T) were acquired at four time points (−7, 30, 90 and 120 dpi) for M83 hemizygous mice injected with PBS or Hu-PFF. [B] Coronal slices of the mouse brain (from posterior to anterior) with t-statistical map overlay; demonstrating the effects of M83 Hu-PFF-induced brain atrophy in terms of voxel-wise volumetric trajectories over time. [C-E] Plots of relative volume change over time (compared to the mean volume per group of interest at the pre-injection time point; −7 dpi) for a peak voxel in [E] the injection site (right dentate gyrus (DG)), and other hippocampal subfields: [C] CA3 and [D] CA1. [F] Spearman correlations between simulated and empirical atrophy derived from M83 Hu-PFF (compared to M83 PBS-injected) mice at −7, 30, 90 and 120 dpi. Correlations are shown as a function of simulation time. After reaching the peak value (r = 0.3297), the model fit slightly drops and stabilizes. [G] Plots of regional pathology (empirical atrophy values at 120 dpi) versus simulated atrophy at the peak model fit (peak correlation timestep); each point represents a region of the Allen Mouse Brain Atlas. [H] Spearman correlation between empirical atrophy at 120 dpi and SNCA expression (r=0.145) and [I] between 120 dpi empirical atrophy and connectivity strength (r=0.167).

Unlike the striatal inoculation which resulted in widespread atrophy, here we observed significant focal volumetric reduction of bilateral hippocampi in M83 Hu-PFF-injected mice. Specifically, we observed significant reductions in volume in bilateral subfields of the hippocampus: dentate gyrus, CA3 and CA1 (Figure 5B-E). Similarly, no mice showed overt motor impairments when examining their behaviour while freely ambulating in their home cages, and no mice succumbed to a humane endpoint beyond the experimental endpoint (>120 dpi). These findings further provide evidence for both regional specificity and connectivity-based spreading of aSyn pathology.

### 2.9. Disease spreading model does not predict atrophy via hippocampal-initiated aSyn spreading

While *in silico* modelling can accurately capture aSyn-induced pathology from striatal seeding of aSyn, it did not accurately predict atrophy from a non-PD-related dentate gyrus seed (Figure 5F-G). Comparisons between simulated and empirical atrophy revealed a peak correlation of r=0.3297 at the 286th time step of the simulation when comparing the atrophy data at 120 dpi between both mouse groups (Figure 5G). Similarly, peak correlations were lower for all other time points (−7 dpi: r=0.0630; 30 dpi: r=0.138; 90 dpi: r=0.215). Notably, when examining the correlation between empirical atrophy at 120 dpi and *SNCA* expression, or compared to connectivity strength measures, neither correlation (r=0.145 and r=0.167, respectively) was sufficiently high enough to capture empirical aSyn spread and its induced pathology using either measure on its own (Figure 5H-I). These simulated findings, along with the empirical data demonstrate that our experiments varying mouse genotype and PFF-species both provide robust atrophy patterns that can be predicted computationally using connectivity and gene expression when injecting into the striatum. These same properties do not extend throughout the brain even though there is evidence of aSyn propagation from a hippocampal epicentre (Bétemps et al. 2014; Dues et al. 2023; Luk et al. 2012b; Rahayel et al. 2022a). These findings suggest that the brain has differential regional vulnerability to aSyn, suggesting that this may need to be further accounted for in the prion-like spreading hypothesis of synucleinopathies.

## 3. Discussion

Our study highlights the intricate interplay between pathogenic aSyn propagation, neuroanatomical and behavioural changes across various mouse models of synucleinopathy. By directly seeding aSyn PFFs into the striatum of WT and hemizygous aSynA53T transgenic mice, we were able to mimic key aspects of human synucleinopathy pathology and investigate the effects of misfolded aSyn spreading on disease susceptibility and progression. Our findings underscore several key points, including the differential effects of aSyn PFF species-dependent inoculation on disease progression, that regional vulnerability is a constraint on neuroanatomical changes induced as evidenced by our striatal and hippocampal PFF inoculations, and that prion-like propagation of aSyn is mediated by this regional vulnerability (which requires further investigation alongside structural connectivity and aSyn gene expression).

Although several studies have examined aSyn spreading in animal models of synucleinopathy via striatal PFF inoculation (Chu et al. 2019; Desplats et al. 2009; Fares et al. 2016; Froula et al. 2019; Luk et al. 2012a; Luk et al. 2012b; Karampetsou et al. 2017; Masuda-Suzukake et al. 2013, 2014; Mougenot et al. 2012; Panattoni et al. 2018; Paumier et al. 2015; Sacino et al. 2014; Tapias et al. 2017; Tullo et al. 2023, 2025; Watts et al. 2013), a full characterization comparing these models between themselves has not yet been explored. Our findings with regards to this comparison are two-fold. First, we observed a dramatic reduction in survival rates for M83 PFF-injected mice. The presence of the A53T mutation in the human *SNCA* gene confers a heightened propensity for aggregation and neurotoxicity (Giasson et al. 2002; Maries et al. 2003), leading to an accelerated disease course compared to the mice only expressing wild type mouse aSyn. This accelerated disease trajectory underscores the heightened vulnerability of possessing the deterministic transgene harboring a mutation that is well-implicated in an autosomal dominant form of PD (Lesage and Brice 2012; Maries et al. 2003). Second, within the M83 PFF inoculated mice, we observed a notable difference in survival rates such that the mice receiving the mouse-derived aSyn fibrils (Ms-PFF) exhibited a more rapid decline along with a more extensive and widespread pattern of atrophy, compared to those injected with human-derived aSyn fibrils (Hu-PFF). This observation suggests a nuanced interaction between the species origin of PFF, the mouse genotype, and disease progression, potentially implicating species-specific factors in modulating aSyn propagation and toxicity. Since the M83 mice also express endogenous mouse aSyn, it is likely that two species-independent processes of aSyn misfolding are occurring: endogenous mouse aSyn misfolding because of the Ms-PFF inoculation, as well as the baseline mutated human aSyn misfolding pathway (as a result of the transgene expression) (Luk et al. 2016; Masuda-Suzukake et al. 2013). However, cross-species templating may also be occurring, albeit at a much lower efficiency (Luk et al. 2016). It is known that aSyn fibrils can adopt distinct conformations or strains, characterized by unique structural features and propagation properties (Danzer et al. 2007; Peelaerts and Baekelandt 2016; Peelaerts et al. 2015; Wang et al. 2014). Species-specific variations in the conformation, stability, and post-translational modifications of aSyn fibrils may influence their ability to propagate and induce neurotoxicity *in vivo* (Cascella et al. 2022; Guo et al. 2013; Lau et al. 2020; Peelaerts and Baekelandt 2016; Tanaka et al. 2006). These differences in fibril properties may influence the rate of aSyn aggregation, seeding efficiency, spreading kinetics, and subsequent neurodegeneration, ultimately impacting the pace of disease progression in these M83 mice (Cascella et al. 2022; Danzer et al. 2007; Peelaerts and Baekelandt 2016; Peelaerts et al. 2015). Another explanation for these findings is the host immune response, and neuroinflammatory cascades triggered by the presence of exogenous aSyn fibrils may differ depending on the species origin of the fibrils. Ms-PFF may induce a stronger inflammatory response in M83 mice due to the greater disparity between endogenous wild type mouse aSyn and exogenous mouse-derived fibrils, compared to the closer resemblance of the mutated A53T human aSyn with exogenous Hu-PFF (Luk et al. 2016). This heightened immune response could contribute to exacerbated neuroinflammation and neuronal dysfunction in M83 mice inoculated with Ms-PFF (Chen et al. 2016; Gao et al. 2011; Kempuraj et al. 2016; Vieira et al. 2015), and accordingly could contribute to the observed differences in disease progression between mouse and human PFF-injected cohorts. Importantly, these findings underscore the importance of considering species-specific effects when modeling synucleinopathy in preclinical studies. However further investigation into the molecular mechanisms underlying these species-specific effects is needed to elucidate such phenotypic differences, and accordingly to provide insights into factors driving aSyn propagation and synucleinopathy pathogenesis.

Regardless of the species of PFF, the neuroanatomical changes were not well predicted by regions connected to the injection site, suggesting that prion-like propagation of aSyn pathology may not be mediated by axonal connectivity. Instead, the combination of structural connectivity and *SNCA* expression levels may better explain the observed pattern of pathology, as evidenced by our *in silico* modeling efforts using a S-I-R agent-based model. While computational models of spread have been explored with respect to aSyn propagation (Rahayel et al. 2022a; Henderson et al. 2019), modeling and simulating aSyn-induced atrophy in mice have yet to be investigated to our knowledge. Our model accurately recapitulated the observed pattern of brain atrophy following striatal PFF seeding, highlighting the potential utility of such models in understanding and predicting synucleinopathy progression. This modeling approach could prove to be a crucial translational tool in bridging the gap between experimental findings from preclinical models and clinical observations, offering insights into the dynamics of aSyn propagation and disease progression in synucleinopathies.

The results of our hippocampal *in vivo* inoculation experiment revealed a region-specific pattern of neuroanatomical changes, characterized by focal volumetric reduction in bilateral hippocampi over time. This finding appears to suggest that previous reports of widespread pathological aSyn deposition from different epicentres (Bétemps et al. 2014; Dues et al. 2023; Luk et al. 2012b; Sacino et al. 2014; Rahayel et al. 2022a) are not sufficient to necessarily relate to downstream atrophy. To ascertain the cellular underpinnings of the observed volumetric changes detected by MRI, and whether they correspond to the deposition of toxic aSyn or other pathological markers, additional immunostaining data is required. Notably, our previous study (Tullo et al. 2023) demonstrated such correspondence following striatal inoculation, suggesting that MRI-derived atrophy data may more likely be influenced by downstream inflammatory processes mediated by microglia and astrocytes, in addition to direct effects of pathological aSyn inclusions in regions connected to the PFF injection site. In fact, in atrophied regions (as identified using MRI), an increased presence of astrocyte and/or microglia markers were observed in some regions that were not always accompanied by increased deposition of phosphorylated aSyn, thereby suggesting an infiltration of atrophy- and inflammation-mediating cells prior to severe phosphorylated aSyn deposition. It is worth investigating whether atrophy primarily results from a toxic neuroinflammatory environment, potentially through refinement of *in silico* models, like the ones used in this study. However, because inflammatory processes are poorly conserved across species compared to humans (Godec et al. 2016), this question remains an open area for future research.

While our study provides valuable insights into the propagation of aSyn pathology and its impact on disease progression, there are important limitations to consider. While our *in vivo* imaging and computational modeling efforts provide valuable insights, they inherently have limitations in capturing the complexity of aSyn propagation and its induced pathology with regards to neuronal loss, particularly in the context of modeling aSyn propagation from a non-dopaminergic region, such as the hippocampus. The limited accuracy of our *in silico* modeling efforts in recapitulating the observed pattern of brain atrophy following hippocampal PFF seeding highlights the challenges associated with extrapolating findings from limited-impacted brain regions to understanding disease spread. There may be nuances related to synucleinopathy-associated pathogenesis that cannot simply be recapitulated using a similarly highly connected brain node. This limitation underscores the need for caution when interpreting results from studies involving these ‘non-traditional’ injection sites and emphasizes the importance of considering regional differences in aSyn propagation dynamics. Notably, *SNCA* expression in the dentate gyrus (in situ hybridization [ISH] signal = 0.989) is actually higher than in the caudoputamen (ISH = 0.525) (Oh et al. 2014), potentially confounding interpretation. Another potential “postive-control” disease epicenter may have been a thalamic nucleus, such as the ventral posteromedial nucleus (VPM), which remains a highly connected hub but expresses lower levels of SNCA (ISH = 0.145) (Oh et al. 2014). It is important to note that the agent-based simulations of aSyn pathogenesis are based on gene expression profiles and structural connectivity data from tracer experiments performed solely on male C57BL/6 mice (Oh et al. 2014). However, these simulations are compared to empirical atrophy data derived from transgenic mice (where the transgene, human A53T SNCA, expression is driven by a prion promoter with its own endogenous expression profile), maintained on a mixed B6/C3H background. Thus, given these differences, there is an inherent limitation of assessing the true accuracy of the *in silico* simulations of empirical aSyn spread and pathology. Future research should aim to address these limitations by refining modeling approaches and integrating data from multiple modalities (including cellular-level gene expression) to provide a more comprehensive understanding of disease pathogenesis. For instance, beyond examining aSyn deposition using immunostaining techniques on postmortem samples, utilizing other imaging techniques, such as positron emission tomography (PET) imaging to examine *in vivo* aSyn aggregates load (as a quantitative measure of “I” agents per region) at different time points, or the integration of functional magnetic resonance imaging (fMRI) data with computational modeling approaches, in which has been previously shown to improve the accuracy of simulated aSyn spreading and pathology in humans, by incorporating activity-dependent spreading via functionally connected regions (Zheng et al. 2019; Rahayel et al. 2023). Finally, investigating the role of regional differences in neuronal connectivity, protein clearance mechanisms, and immune responses in modulating the spread of aSyn pathology may provide valuable insights into the pathophysiology of synucleinopathies and identify novel therapeutic targets.

In summary, our study provides novel insights into the complex interplay between genetic predisposition, aSyn species, and disease epicenter in driving synucleinopathy pathogenesis and progression. These findings have implications for the development of targeted therapeutic interventions aimed at halting or slowing disease progression in PD and other synucleinopathies. However, further research is warranted to elucidate the underlying mechanisms driving aSyn propagation and to refine predictive models of disease progression.

## 4. Methods

### 4.1. Animals

We used transgenic hemizygous M83 mice (B6; C3H-Tg[SNCA]83Vle/J), expressing one copy of human aSyn bearing the familial PD-related A53T mutation under the control of the mouse prion protein promoter (TgM83+/-) (Giasson et al. 2002), maintained on a C57BL6/C3H background. The M83 hemizygous mice used here present no phenotype until they are 22 and 28 months of age, on average (Giasson et al. 2002), unless aSyn PFFs are injected to trigger accumulation of toxic aSyn. Wild-type (WT) mice, littermates of the M83 mice, have no copies of the aSyn transgene and are maintained on the same mouse line (C57BL/6 x C3H). All mice were bred in-house via the following breeding scheme: TgM83+/- x TgM83+/-, where 50% of the pups were TgM83+/- (i.e. M83 hemizygous) and 25% of the pups were TgM83-/- (i.e. WT).

All mice were housed at the Douglas Research Center Animal Facility (McGill University, Montreal, QC, Canada) under standard housing conditions with food and water ad libitum. Mice were typically housed in maximum groups of four, and mice were exceptionally singly housed when fighting occurred (only seen in a small proportion of males). Mice were housed with a 12/12-hour light/dark cycle, with lights on at 08:00. All study procedures were performed in accordance with the Canadian Council on Animal Care and approved by the McGill University Animal Care Committee and the Douglas Facility Animal Care Committee under the animal use protocol (AUP DOUG-8068).

The experimental timeline with all the procedures performed at each time point is depicted in Figure 1A. Prior to any intervention, the mice underwent weighing: prior to surgery and at each time point prior to MRI, as well as prior to the experimental or humane endpoint, before perfusion. Final numbers per group per experimental condition per time point are presented in Supplementary Table 1.

### 4.2. Injection material and Stereotaxic injections

To investigate toxic aSyn spreading, we injected healthy 3-4 month old mice with preformed fibrils (PFF) of human (Hu-) or mouse (Ms-) aSyn, or phosphate-buffered saline (PBS) in the right caudoputamen (dorsal striatum) to have a known disease epicentre of spreading. Injection group assignment was random, with an equal number of mice across injection assignment, genotype, and sex.

Hu-PFF and Ms-PFF were generated in-house (https://zenodo.org/record/6430401#.YrsWKJPMK8X) and were sonicated using a Bioruptor Pico sonication device (Diagenode, USA) preset at 5°C water bath for at least 60 cycles of 30 seconds on and 30 seconds off. PFF were characterized using a negative staining (https://zenodo.org/record/6524616#.YrsWJpPMK8V) and were visualized using a transmission electron microscope (Tecnai G2 Spirit Twin 120 kV Cryo-TEM) coupled to a camera (Gatan Ultrascan 4000 4 k × 4 k CCD Camera model 895) and analyzed with Fiji-ImageJ1.5 and GraphPad Prism 9 software. PFF were characterized by dynamic light scattering (DLS) in addition to electron microscopy (EM). EM characterization of the fibrils in terms of distribution per length can be seen in Supplementary Figure 1.

Briefly, the mice were anaesthetized with isoflurane (5% induction, 2% maintenance), given an injection of carprofen (0.1cc/10 grams) and xylocaine was applied on the scalp prior to cutting for pain analgesia management. Next, the mice were positioned into a stereotaxic platform. PFFs (total volume: 2.5 μL; 5 mg/mL; total protein concentration, 12.5 µg per brain) were stereotaxically injected in the right dorsal striatum (co-ordinates: +0.2 mm relative to Bregma, +2.0 mm from midline, +2.6 mm beneath the dura) (Luk et al. 2012a; Luk et al. 2012b) or the right hippocampus (co-ordinates: −2.5 mm relative to Bregma, +2.0 mm from midline, +1.8 mm beneath the dura) (Luk et al. 2012a; Iba et al. 2013). Injections were performed using a 5 μL syringe (Hamilton; 33 gauge) at a rate of 0.25 μL per min with the needle in place for about 5 min prior to and after infusion of the inoculum. Different syringes and needles were used for each type of inoculum to prevent any cross-contamination. Post-injection, the mice were placed on a heating pad for recovery before being returned to their home cage. Post-injection experimenters were blind to the injection assignment (as well as genotype) of each mouse during the data acquisition processes (MRI and behavioural testing), as well as for the analysis.

After inoculation, the mice were monitored weekly for health and neurological signs such as reduced grooming, kyphosis, and/or decreased motor functioning (including reduced ambulation, tail rigidity, paraparesis), whereby the frequency of monitoring increased to daily checks upon the anticipation of the onset of motor symptomatology (∼80 dpi; 10 days prior to the typical onset of motor impairments; (Froula et al. 2019; Luk et al. 2012a; Luk et al. 2012b; Masuda-Suzukake et al. 2013, 2014; Mougenot et al. 2012; Sacino et al. 2014; Tullo et al. 2023, 2025; Watts et al. 2013)), up until the experimental or humane endpoint.

### 4.3. Magnetic Resonance Imaging (MRI) acquisition

Mice underwent longitudinal MRI at four time points: −7, 30, 90 and 120 days post-injection, unless they reached a humane endpoint prior to any experimental timepoint. MRI acquisition was performed on 7.0-T Bruker Biospec 70/30 USR 30-cm inner bore diameter (with Bruker’s MRI CryoProbe and AVANCE NEO electronics) at the Douglas Research Centre (Montreal, QC, Canada). High-resolution *in vivo* T1-weighted images (FLASH; Fast Low Angle SHot) were acquired for each subject (TE/TR of 4.5 ms/20 ms, 100 μm^3^ isotropic voxels, 2 averages, scan time= 14 min, flip angle=20°). For each time point acquisition, the mice were anaesthetized with isoflurane at 3% induction (with 1% oxygen flow rate) for 3:30 minutes whereby a bolus injection of dexmedetomidine (1:240 dilution) was administered intraperitoneally. The mice remained in the induction chamber until 5:30 minutes elapsed by which the mice were transferred to the MRI scanner and placed under 1.5% isoflurane with a constant infusion of dexmedetomidine (0.05 mg/kg/hr).

#### 4.3.1. Image processing

##### 4.3.1.1. Pre-processing

All brain images were exported as DICOM from the scanner and converted to the MINC (Medical Imaging NetCDF) file format. Image processing was performed using the MINC suite of software tools (http://bic-mni.github.io). At this stage, the images were manually inspected for artifacts (hardware, software, motion artifacts or tissue heterogeneity and foreign bodies) and images with said artifacts would be excluded (https://github.com/CoBrALab/documentation/wiki/Mouse-QC-Manual-(Structural)).

After passing quality control, the images were stripped of their native coordinate system, left-right flipped to compensate for Bruker’s native radiological coordinate system, denoised using patch-based adaptive non-local means algorithm (Coupe et al. 2008), and affinely registered to an average mouse template (the Dorr–Steadman–Ullman atlas; (Dorr et al. 2008; Steadman et al. 2014; Ullmann et al. 2014)) to produce a rough brain mask. Next, a bias field correction was performed, and intensity inhomogeneity was corrected using N4ITK (Tustison et al., 2010) to a minimum distance of 5 mm. We employed a standard operating procedure as previously implemented as by our group (Gallino et al. 2019; Guma et al. 2021; Rollins et al. 2019; Tullo et al. 2023, 2025) (https://github.com/CobraLab/documentation/wiki/Mouse-Scan-Preprocessing).

##### 4.3.1.2. Deformation-based morphometry

The denoised and inhomogeneity-corrected images were aligned through a series of linear and nonlinear registration steps with iterative template refinement using the ANTS registration algorithm (Avants et al. 2011) to generate a study-specific template enabling voxel correspondence for every brain voxel across every subject time point image. Given the longitudinal nature of the data, a two-level deformation-based morphometry technique ((Germann et al. 2025); antsMultivariateTemplateConstruction2.sh; https://github.com/CoBrALab/twolevel_ants_dbm) was used such that all time point images per subject were first registered to generate a subject-wise template (first level), followed by the registration of all the subject-wise templates to create a final group-wise template (second level). The minimum deformation required at a voxel-level to map each subject time point image to its subject template was log-transformed and blurred with Gaussian smoothing using ∼0.085 mm full width at half maximum kernel to better conform to normative distribution assumptions for statistical testing. Voxel-wise volume measures were derived from the relative Jacobian determinants to measure local neuroanatomical change over time (exclusively modelling the nonlinear transformations of the deformations) (Chung et al. 2001). These Jacobian determinants can be either expansions or contractions and are dependent on the magnitude of the deformation at each voxel. The outputs were inspected to assess quality control of the registration by visually assessing the resultant images of the registrations for a proper orientation, size, and sensible group average, prior to performing the analyses. This method has been previously used by our group to examine voxel-wise volumetric change over time (Gallino et al. 2019; Guma et al. 2021; Kong et al. 2018; Rollins et al. 2019; Tullo et al. 2023, 2025).

### 4.4. Motor behavioural tasks

On the subsequent days after the MRI scanning day (>24 hours post-scanning) at each of the four timepoints (−7, 30, 90, 120 dpi), the mice underwent behavioural testing with one behavioural test administered each day. The following tasks were performed as follows: pole test, rotarod and wire hang. The behavioural tests were conducted during the light phase of the day cycle, between 8 am and 8 pm for each test. The mice were habituated to the testing room for one hour prior to administering the behavioural test.

#### 4.4.1. Pole Test

The pole test was administered to examine motor agility and fine motor movements (Matsuura et al. 1997). Each mouse was placed head-upward on the top of a vertical rough-surfaced pole (wooden dowel covered in surgical tape; diameter 8 mm; height 55 cm). The time required for the mouse to descend to the floor was recorded with a maximum duration of 2 minutes (120 seconds). Each subject completed three trials with a minimum 30-minute rest between each trial, following the protocol detailed by Matsuura et al. (1997). Notably, two scores were recorded: 1) whether the task was performed successfully or was a failed attempt, and 2) the time taken to descend. If a mouse was not able to turn downward and remained at the top of the pole, the trial was marked as a failure and the maximum time allotted was noted (120 seconds). In cases where the mouse fell part of the way down but subsequently descended the rest, the behaviour was scored as a failure however the time it took to reach the floor was still noted. If the mouse fell for more than half the length of the pole, the behaviour was scored as a failure and the maximum time allotted was noted. For cases where a mouse fell immediately after placement at the top of the pole, a failed attempt, the trial was repeated.

#### 4.4.2. Rotarod

To investigate motor coordination and balance, mice were tested on the Rotarod as previously described (Janickova et al. 2017). Mice were placed on the Rotarod (San Diego Instruments; San Diego, CA, USA) and rotation was accelerated linearly from 4 to 25 revolutions per minute (rpm), increasing rpm every 15 seconds. Time spent walking on top of the rod before falling off the rod or hanging on and riding completely around the rod was recorded. Mice were given three trials with a minimum 30-minute inter-trial rest interval.

#### 4.4.3. Wire-Hang

Neuromuscular strength was tested with the wire hang test. The mouse was placed on a wire by waving it gently so that it gripped the wire and then inverted. Latency to fall over the course of three trials was recorded with a 3-minute cut-off time, with a minimum 30-minute inter-trial rest period (Froula et al. 2019). Similar to the pole test, performance on the wire hang was examined in terms of latency and failure/success rates. If the mouse holds on for more than 3 minutes, it receives the maximum allotted time (180 seconds) and a successful score. For this task, there are three possible outcomes: 1) the mouse holds on for more than 3 minutes, 2) the mouse holds on for less than 3 minutes and receives a failure score with the duration recorded, or in the most extreme case, 3) the mouse is unable to hold on for at least 10 seconds. In the latter case, three attempts were allowed during the trial before a failure score and a duration of 0 seconds was recorded.

### 4.5 Statistical analyses

Statistical analyses were carried out using R software (3.5.0) with the survival package (https://cran.r-project.org/web/packages/survival/index.html) for the Cox proportional hazards modelling and the RMINC package (https://github.com/Mouse-Imaging-Centre/RMINC) for voxel-wise linear regression. No data points were excluded for any statistical analysis.

#### 4.5.1. Behavioural analysis

Disease progression (survival analysis) was examined using Cox Proportional Hazard modeling (Kassambara 2013; Tullo et al. 2025). This cumulative incidence model is considered superior to the t-test or ANOVA given the nature of the data where two measures can be used to denote the behaviour of the mouse. For the survival analysis, we examined 1) the average dpi to succumb to disease (reach a humane endpoint) as well as 2) the proportion of the mice that succumbed at that dpi. Sex was included in the analysis based on sex differences in disease progression previously observed (Tullo et al. 2025).

Differences in onset of overt motor symptoms (first observed in ambulating mice in terms of dpi) as well as motor symptom duration (endpoint dpi minus onset dpi) was assessed using general linear models. Motor symptom onset was examined for the three way interaction of genotype by injection group by sex (average dpi ∼ genotype*injection_group*sex). For motor symptom duration, given that no WT mice nor female M83 Hu-PFF injected mice succumbed to a humane endpoint prior to the experimental time point, the main effect of the injection group was examined (duration ∼ injection_group). Post-hoc testing was performed using Tukey HSD for multiple comparisons corrections.

For longitudinal measures such as weight and rotarod performance (averaged across trials at each time point), linear mixed effect models were used to assess genotype by injection group by sex differences across the time points (genotype*injection_group*sex*time point + (1|subject_ID)). For weight and rotarod, the addition of a random effect of cohort and litter size was added to the model, as determined by Akaike information criterion (AIC) (Akaike 1974) separately. For pole test and wire hang tests, performance was examined with regards to latency and failure/success rate using a Cox Proportional Hazard model (Tullo et al. 2025) as these two tasks have strict time cut-offs to determine the success/fail rate. Distribution of data, highlighting a ceiling and floor effect for wire hang and pole test respectively, is available in the Supplementary Figures 10 and 11, as well as genotype by injection group differences at timepoints −7 and 30 dpi, and repeated analyses at each timepoint taking into account sex differences (Supplementary Figure 3).

#### 4.5.2. Longitudinal volumetric analysis

We examined longitudinal volume trajectories to investigate subject-wise changes over time based on the injection, genotype and sex of the mice. Mixed linear models were performed at each voxel:

Relative volume at each voxel ∼ genotype*injection_group*time point + sex + (1| subject_ID) as determined by AIC. The False Discovery Rate (FDR) (Benjamini and Hochberg 1995) correction was applied to control for multiple comparisons.

#### 4.5.3. Partial least squares analysis

Partial Least Squares (PLS) is a statistical method used for multivariate analysis to determine the optimal weighted linear combination of two sets of variables: whole brain voxel-wise DBM and z-scored behavioural and demographic metrics, aiming to maximize their covariance (Zeighami et al. 2019; McIntosh and Mišić 2013; McIntosh and Lobaugh 2004). Here, PLS was employed to investigate the relationship between MRI-derived atrophy (voxel-wise relative Jacobian difference between the −7 & 90 dpi symptom onset time point) and symptom profile (demographic and symptom-based variables such as sex, weight, genotype, injection, symptom score, and motor performance on the pole test, rotarod and wire hang motor tasks).

PLS results in a series of orthogonal latent variables (LV), representing ‘brain weights’ and ‘behaviour weights’, indicating how each voxel or behavioural variable contributes to a given LV. Additionally, it produces a singular value indicating the proportion of covariance explained by the LV (Eckart and Young 1936). To evaluate the statistical significance and the contribution of original variables to each LV, permutation testing (n=1000) and bootstrap resampling (n=1000) were conducted.

Permutation testing was utilized to evaluate the statistical significance of each LV by randomly shuffling the rows (subjects) of the brain data matrix to nullify dependencies between brain and behaviour (n=1000 repetitions) (Guma et al. 2021). This process generated a null distribution of possible brain-behaviour correlations, and singular value decomposition (SVD) was applied to these “null” correlations to derive a distribution of singular values under the null hypothesis. The probability that a permuted singular value exceeds the original singular value allows for the calculation of a p-value (Zeighami et al. 2019; Patel et al. 2020), with a threshold of p<0.05 considered statistically significant (indicating a 95% or greater chance that the singular value of the non-permuted data exceeds that of a permuted singular value).

Bootstrap resampling was employed to evaluate the contribution of individual brain and behaviour variables to each LV. Subjects (rows for both matrices) were randomly sampled and replaced (n=1000), generating a set of resampled correlation matrices to which SVD was applied, creating a sampling distribution for each weight of the singular vectors (McIntosh and Mišić 2013). The ratio of each singular vector weight to its bootstrap-estimated standard error was calculated as a “bootstrap ratio” (BSR) for each voxel. This analysis allows for the identification of voxels making substantial contributions to specific patterns based on large bootstrap ratios. We used a BSR threshold of 2.58, analogous to a p-value of 0.01 (Krishnan et al. 2011; McIntosh and Lobaugh 2004; Nordin et al. 2018; Persson et al. 2014; Zeighami et al. 2019).

#### 4.5.4. Anatomical connectivity and SNCA gene expression

Data from the Allen Mouse Brain Connectivity Atlas (AMBCA) (Oh et al. 2014; Kuan et al. 2015) was used to examine whether PFF-induced atrophy was correlated with structural connectivity projecting from the disease epicenter (our injection site in the experimental mice). The AMBCA is a high-resolution map of mouse whole brain neuronal connections. The atlas consists of high-resolution images of axonal projections derived using enhanced green fluorescent protein (EGFP)-expressing adeno-associated viral vectors to trace axonal projections from defined regions and cell types, derived from both WT mice and Cre-driver mice, and using serial two-photon tomography (Oh et al. 2014; Kuan et al. 2015). The connectivity matrix consisted of 213 rows representing source regions and 426 columns representing the 213 ipsilateral regions and the 213 contralateral regions. Each element in the matrix represents the tracer-derived connectivity strength of every incoming connection from a region (injection site; source region) (Oh et al. 2014).

Gene expression data for each of the 426 parcellated Allen regions was obtained from the Allen Mouse Brain Atlas API (http://help.brain-map.org/display/mousebrain/API) (Lein et al. 2007; Ng et al. 2009). The section images containing gene expression values were divided into a 200 × 200-μm grid to generate a low-resolution three-dimensional volume and then mapped to the three-dimensional reference model. Pixel-based gene expression statistics were then summarized within each division and expression energy was chosen as the gene expression measurement. For each experiment, the expression energy of each region was calculated by unionizing the corresponding grid voxels with the same structure label in the atlas. These values were subsequently z-scored and averaged across the corresponding experiments for each region (Rahayel et al. 2022a).

### 4.6. S-I-R model

Here, we used the Susceptible-Infected-Removed (S-I-R) agent-based model to explore temporal spreading of pathological aSyn and quantify the atrophy induced based on the existing code from Zheng et al. (2019) and Rahayel et al. (2022a; 2022b). While there are different approaches for modeling epidemic spread over a network, most commonly diffusion models, where propagation is modeled as a concentration, obeying the law of mass effect with first-order kinetics (Henderson et al. 2019; Zheng et al. 2019), agent-based models allow for explanatory power such that simple local interactions can translate into complex global behaviour (Zheng et al. 2019). For example, the infectious state of each individual agent and its motility are simulated, and biological properties of the system can be incorporated to simulate the spread, such as regions with high infectious protein levels would likely have reduced connectivity and accordingly limited ability to further propagate (Zheng et al. 2019).

To model the propagation, aSyn proteins will be coded as agents in one of three states, “Susceptible” (normal) state, “Infectious” (abnormal misfolded form), or “Removed” (metabolized protein); thus, being referred to as a S-I-R model. As agents move from one brain region to another along the brain’s connectome, via projections derived from the AMBA connectivity atlas (Oh et al. 2014), contact between “I” and “S” proteins transforms the “S” into “I”, according to a certain probability, and the accumulation of “I” proteins over time leads to degeneration and cell death within that region (Zheng et al. 2019). While normal aSyn “S” is continuously synthesised throughout the brain, as informed by *SNCA* gene expression data obtained from the Allen Mouse Brain Gene Expression Atlas (Lein et al. 2007; Ng et al. 2009), misfolded aSyn “I” will be initially seeded in the injection site based on the experimental mice data. Assessing model performance was achieved by examining differences between actual patterns of atrophy in the mice against simulated measures of pathogenic aSyn-induced pathology.

An atrophy map was generated from the DBM data, such that for each region defined in the AMBA atlas, an average of the unsmoothed, linear and nonlinear (including the residual global linear transformations attributable to differences in total brain size for example) Jacobian determinants at each voxel was computed for each of the 426 AMBA regions (Zheng et al. 2019). Next, the atrophy value was computed as a t-statistic describing the difference between the most symptomatic group (M83 Ms-PFF) relative to the control group (WT PBS). This process was repeated for the data from hippocampal injection experiments; however here the comparison was between the M83 Hu-PFF and the M83 PBS-injected mice.

### 4.7. Data availability

All source data (demographic, behavioural, atrophy and simulated data) has been made publicly available in an online repository (doi.org/10.5281/zenodo.16614413). Any other information is available upon request.

### 4.8. Code availability

All our methods and techniques (preprocessing pipeline, DBM, SIR, etc.) used in this manuscript are open source and available at our Github (https://github.com/CoBrALab) or available on an online repository (doi.org/10.5281/zenodo.16614413).

### 4.10. Author Contributions

S.T. with the help of M.M.C. conceived, designed, and planned the experiments. S.T. along with J.S.H.P, D.G., M.P., and K.M. collected the MRI and behavioural data. W.L., E.D.C.P., and I.S. produced and characterized the PFF inoculum, under the supervision of T.M.D. and E.A.F. S.T. processed and analyzed the MRI and behavioural data. J.S.H.P. performed the experiment with the hippocampal inoculation. S.T. adapted the SIR model from existing code developed by S.R. and Y.Q.Z. with help from B.M. and A.D. S.T. performed the computational modeling analysis. SIR interpretation was performed with the help of S.R. and A.D. S.T. wrote the manuscript with support from M.M.C.

