## Supplementary Information for "In vivo and in silico alpha-synuclein propagation dynamics: The role of genotype, epicentre, and connectivity"

| | | -7 & 30 dpi | | | | 90 dpi | | | | 120 dpi | | | |
|---|---|---|---|---|---|---|---|---|---|---|---|---|---|
| | | PBS | Hu-PFF | Ms-PFF | Total | PBS | Hu-PFF | Ms-PFF | Total | PBS | Hu-PFF | Ms-PFF | Total |
| WT | M | 28 | 26 | 28 | **82** | 20 | 19 | 15 | **54** | 8 | 8 | 9 | **25** |
| | F | 28 | 29 | 26 | **83** | 17 | 19 | 17 | **53** | 8 | 9 | 7 | **24** |
| M83 | M | 30 | 34 | 32 | **96** | 21 | 23 | 23 | **67** | 9 | 5 | 6 | **20** |
| | F | 31 | 33 | 33 | **97** | 23 | 20 | 23 | **66** | 9 | 8 | 3 | **20** |
| | | | | | **358** | | | | **240** | | | | **89** |

**Supplementary Table 1. Number of mice per time point for MRI and behavioural testing.** Wild-type (WT); M83 hemizygous aSynA53T transgenic mice (M83); phosphate buffered saline (PBS); human aSyn preformed fibrils (Hu-PFF); mouse aSyn preformed fibrils (Ms-PFF); male (M); female (F); days post-injection (dpi).

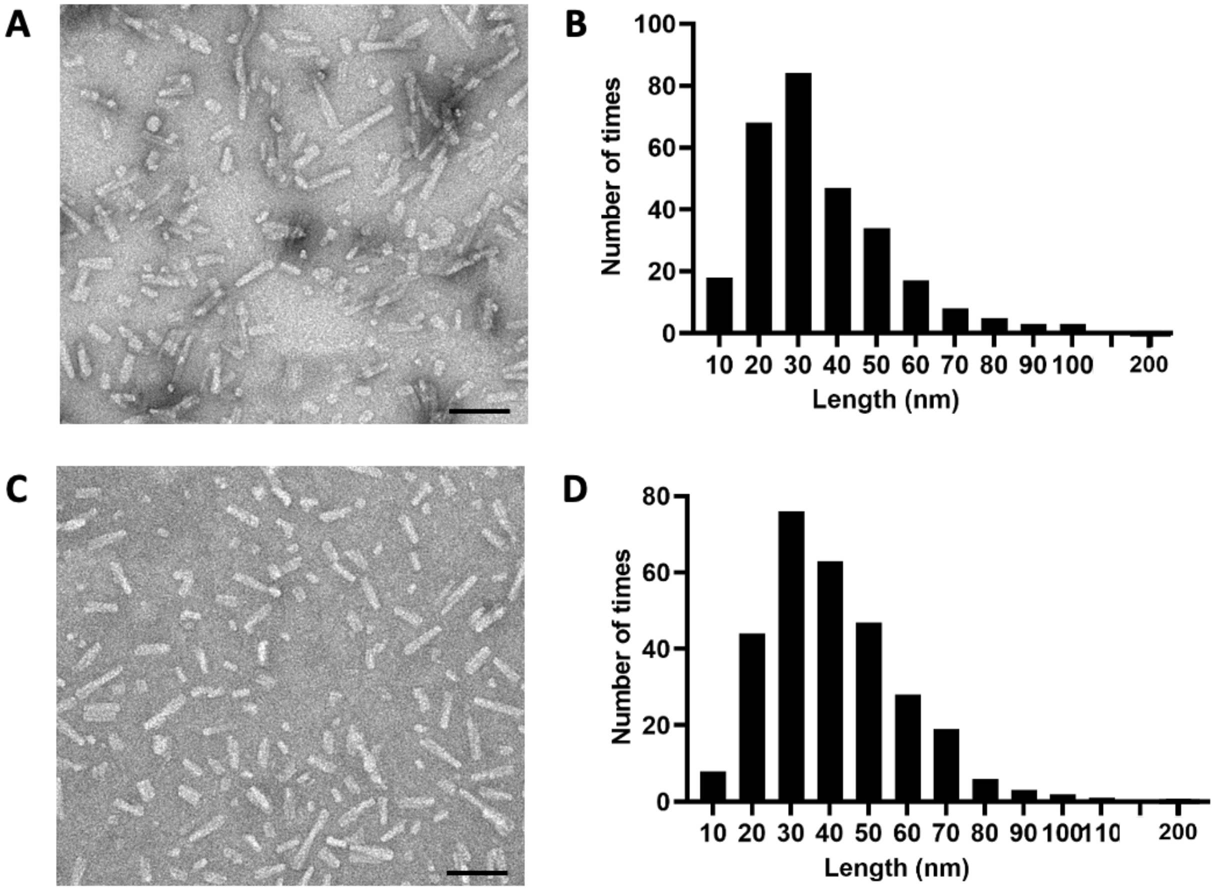


**Supplementary Figure 1. Human and mouse alpha-synuclein preformed fibrils characterization.** [A,C] Representative photomicrographs of human (Hu-) [A] and mouse (Ms-)

[C] aSyn PFF stained by negative staining and visualized using Tecnai G2 Spirit electron microscope. Bar scale 100 nm. [B,D] Histograms show PFF length distribution measured using ImageJ software and their distribution plotted using GraphPad Prism software. [B] Hu-PFF were sonicated for 60 cycles of 30-s On/30-s Off (n= 288, length average= 35.76 nm, median length= 31.65 nm, minimal length = 9 nm, maximal length= 200 nm). [D] Ms-PFF were sonicated for 60 cycles of 30-s On/30-s Off (n= 298, length average= 41.48 nm, median length= 37.45 nm, minimal length = 9 nm, maximal length= 200.6 nm).

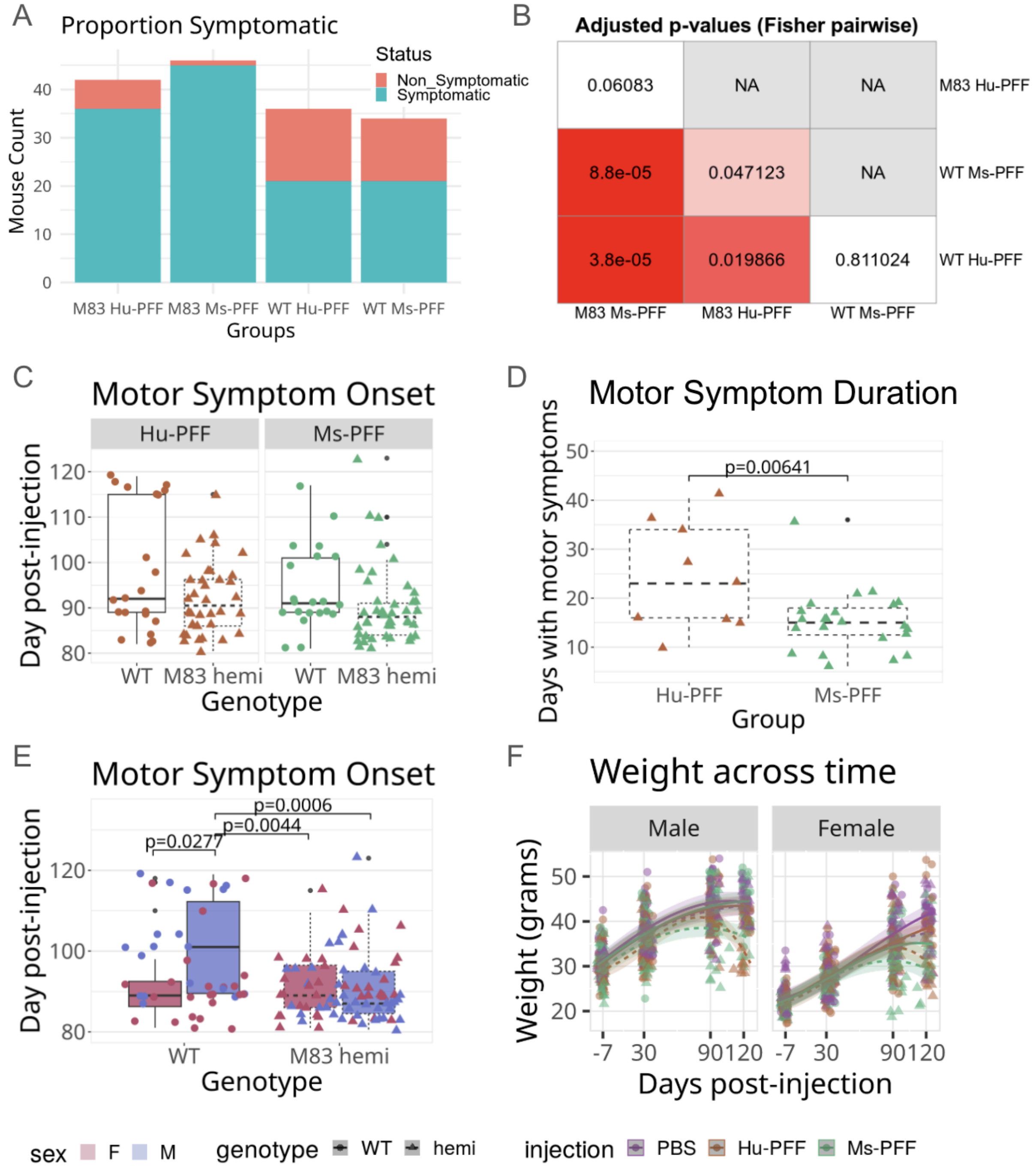


**Supplementary Figure 2. Overt motor symptom onset and disease progression measures (symptom duration and weight) reveal sex differences within PFF-injected mice.**

[A] Proportion of symptomatic mice grouped by genotype and PFF species inoculum; M83 Hu-PFF: n=36/42; M83 Ms-PFF: n=45/46; WT Hu-PFF: n=21/36; WT Ms-PFF: n=21/34. [B] Fisher's Exact Test for Count Data pairwise comparisons between each of the four groups were performed. The significant difference between M83 Ms-PFF and M83 Hu-PFF (p=0.0507) did not survive multiple comparisons correction (p=0.0603). Other significant pairwise comparisons after FDR correction were as follows: M83 Ms-PFF vs WT Ms-PFF: p=0.0000292; M83 Ms-PFF vs WT Hu-PFF: p=0.00000631; M83 Hu-PFF vs WT Ms-PFF: p=0.0314; M83 Hu-PFF vs WT Hu-PFF: p=0.00993. [C] Onset of motor symptoms in terms of the first day of observed symptoms showed no significant differences between PFF-injected groups regardless of mouse genotype. [D] While no differences in overt motor impairment onset between the two PFF-injection groups was observed, we sought to determine if there were differences in the duration of symptoms between these groups given the differences observed in survival. We observed a significant effect of injection (p=0.00641) within the humane endpoint mice (n=34), whereby M83 Ms-PFF-injected mice (n=23; 13 females and 10 males) had significantly shorter durations of visible motor impairment compared to their Hu-PFF injected counterparts (n=9 male mice). [E] Regardless of the species of PFF inoculum, we observed sex differences between the genotypes (p=0.00441); no significant genotype by injection by sex effect, nor a genotype by injection effect was observed. Male WT PFF-injected mice had a significantly later symptom onset compared to female WT PFF-injected mice (p=0.0277), as well as their M83 PFF-injected counterparts (males: p=0.0006; females: p=0.0044). [F] Given the known differences in weight between male and female mice, this relationship was further examined in a sex-specific manner, with similar weight trends between the groups (male: cubic M83 Hu-PFF: df=471.145; p=0.000893; quadratic M83 Ms-PFF: df=485.042; p=0.000181; female: cubic M83 Hu-PFF: df=479.229; p=0.0454; compared to WT PBS mice). This difference was also observed when examining this relationship within each sex between M83 PFF-injected mice: (M83 Hu-PFF vs M83 Ms-PFF for males: df=472; p=0.0027; females: df=482; p=0.0538).

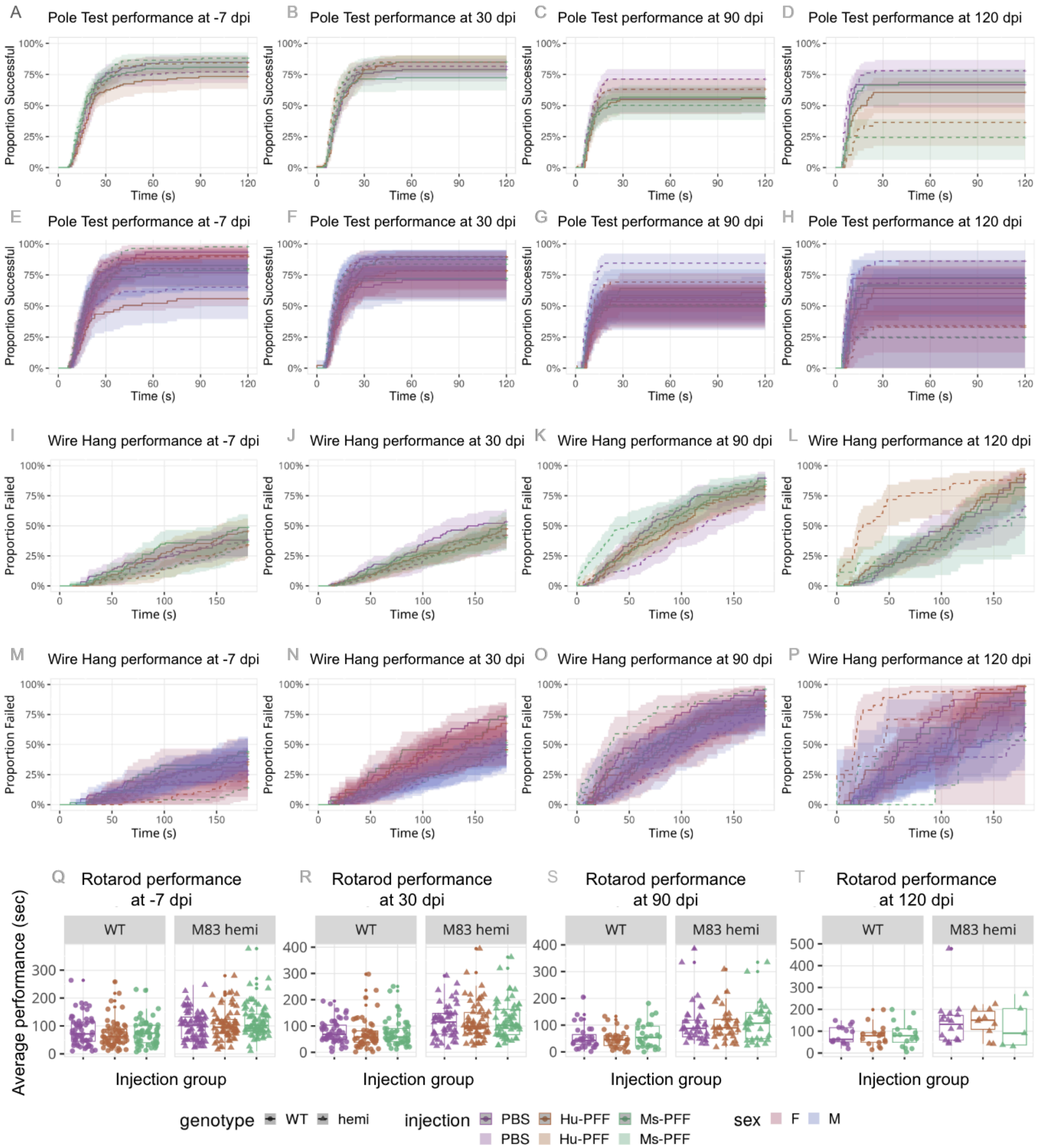


**Supplementary Figure 3. Cross-sectional behavioural analysis with sex differences for the pole test, wire hang and rotarod test at each timepoint.** [A-D] Pole test performance (averaged across trials) at -7, 30, 90, and 120 dpi, examining genotype by injection group differences. Significant genotype by injection interaction effect observed at 90 (M83 Ms-PFF p=0.049881) and 120 dpi (M83 Ms-PFF p= 0.000749; M83 Hu-PFF p=0.00820). Shading represents a 95% confidence interval. [E-H] Sex-specific pole test performance at -7, 30, 90, and

120 dpi, examining genotype by injection group by sex differences. When examining performance differences with regards to genotype and injection within the sexes, at 120 dpi, M83 Ms-PFF mice had worse performances (higher rates of failures and longer descend durations) compared to WT PBS (p=0.000951). Shading represents a 95% confidence interval. [I-L] Wire hang test performance at -7, 30, 90, and 120 dpi, examining genotype by injection group differences. Significant genotype by injection interaction effect observed at 90 (M83 Ms-PFF p=0.000789; M83 Hu-PFF p=0.00691) and 120 dpi (M83 Hu-PFF p=0.000151). Shading represents a 95% confidence interval. [M-P] Sex-specific wire hang test performance at -7, 30, 90, and 120 dpi, examining genotype by injection group by sex differences. Significant sex by genotype by injection interaction effect observed at 90 (female M83 Ms-PFF p=0.049881). When examining performance differences with regards to genotype and injection within the sexes, at 90 and 120 dpi, for both male and female mice, M83 Hu-PFF mice performed worse (higher rates of failures and shorter durations) compared to WT PBS mice (90 dpi: males: p=0.0322; females: p=0.0377; 120 dpi: males: p=0.000203; females: p=0.00697). Furthermore, female M83 Ms-PFF mice had worse performances compared to WT PBS mice at 90 dpi (p=6.97e-5). Shading represents a 95% confidence interval. [Q-T] Average rotarod duration at -7, 30, 90, and 120 dpi, examining genotype by injection group by sex differences. No statistical sex by genotype by injection interaction were observed at each timepoint.When examining performance differences with regards to genotype and injection within the sexes, for the males mice at 120 dpi, we observe longer durations for M83 hemizygous Hu-PFF injected mice compared to their WT counterparts (p=0.0216) and compared to WT PBS-injected mice (p=0.0124). Purple colour denotes PBS-injected mice, orange colour denotes Hu-PFF injected mice, and green colour denotes Ms-PFF injected mice. Line type and data point shapes were used to denote the genotypes: solid line with circular points for WT mice and dashed line with triangular points for M83 hemi mice. When applicable, shading of lines and data points are coloured across to sex; blue for males and red for females.

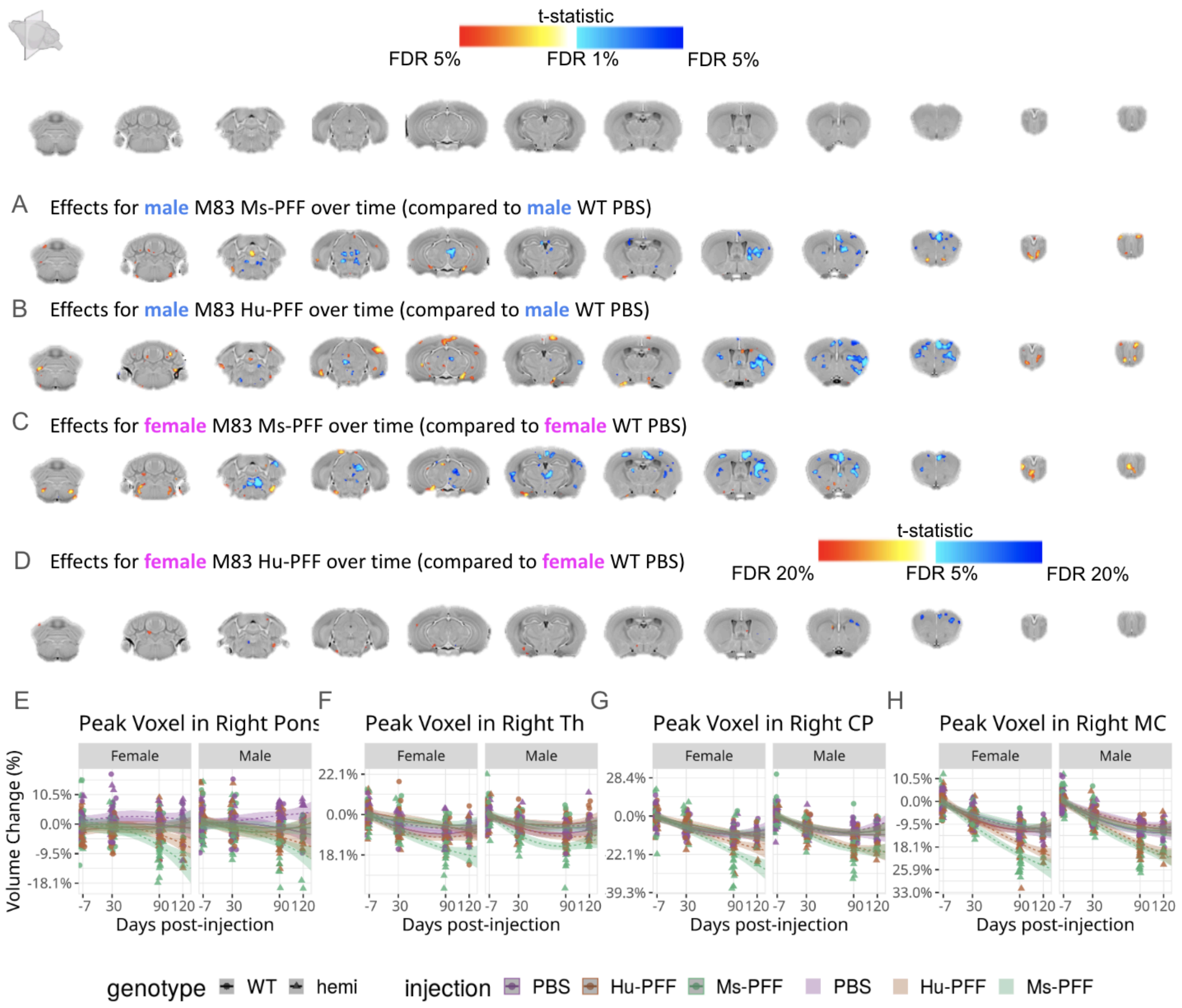


**Supplementary Figure 4. Sex-focused analysis of genotype by injection group differences in voxel-wise volumetric trajectories.** [A-D] Coronal slices of the mouse brain (from posterior to anterior) with t-statistical map overlay; demonstrating the effects for male [A] M83 Ms-PFF-injected and [B] M83 Hu-PFF-injected over time (compared to male WT PBS injected mice), and for the female [C] Ms-PFF-injected and [D] Hu-PFF-injected over time (compared to female WT PBS injected mice). Colour map describes the direction of the t-statistics; cooler colours denoting negative slopes; most commonly corresponding to volume decline and warmer colours denoting positive slopes, corresponding to volume increases over time for the group of interest indicated; t-values bounded between FDR 1 and 5% for maps (A-C) and between FDR 5 and 20% for map (D). [E-H] Plot of percent relative volume change over the four time points (-7, 30, 90 and 120 days post-injection). Shading represents ±1 standard error of the mean. We chose to highlight voxels within regions where group differences over time were observed across comparisons; peak voxel in [E] the right side of the pons, [F] right thalamus (Th), [G] the injection site (right caudoputamen (CP)), [H] right primary motor area (MC). This figure compliments Figure 3 in the main text [A-I], here highlighting the same analysis with respect to each sex. Purple line for PBS-injected mice, orange line for Hu-PFF-injected mice, green line for Ms-PFF-injected mice, solid line and triangle points for WT and dashed line and circular points for M83 hemizygous mice. Steepest rates of decline are observed for Ms-PFF-injected mice regardless of sex.

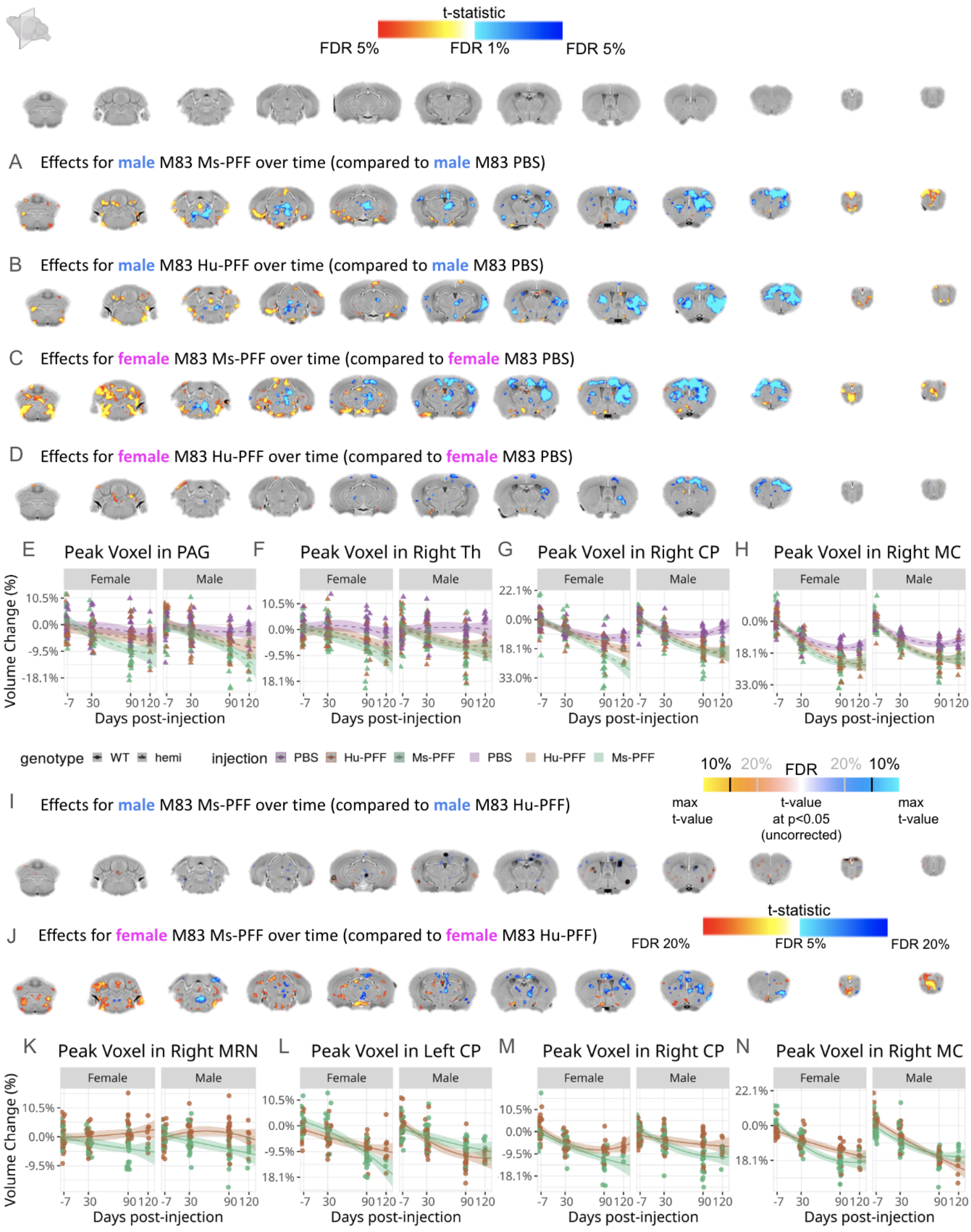

t-statistic
FDR 5%
FDR 1%
FDR 5%
A Effects for male M83 Ms-PFF over time (compared to male M83 PBS)
B Effects for male M83 Hu-PFF over time (compared to male M83 PBS)
C Effects for female M83 Ms-PFF over time (compared to female M83 PBS)
D Effects for female M83 Hu-PFF over time (compared to female M83 PBS)
E Peak Voxel in PAG
F Peak Voxel in Right Th
G Peak Voxel in Right CP
H Peak Voxel in Right MC
Female
Male
Volume Change (%)
Days post-injection
genotype WT hemi
injection PBS Hu-PFF Ms-PFF PBS Hu-PFF Ms-PFF
10% 20% FDR 20% 10%
max t-value
t-value at p<0.05 (uncorrected)
max t-value
I Effects for male M83 Ms-PFF over time (compared to male M83 Hu-PFF)
J Effects for female M83 Ms-PFF over time (compared to female M83 Hu-PFF)
t-statistic
FDR 20%
FDR 5%
FDR 20%
K Peak Voxel in Right MRN
L Peak Voxel in Left CP
M Peak Voxel in Right CP
N Peak Voxel in Right MC
Female
Male
Volume Change (%)
Days post-injection

**Supplementary Figure 5. Genotype (M83) group-focused analysis (with sex-based analysis) of injection group differences in voxel-wise volumetric trajectories.** [A-D; I-J] Coronal slices of the mouse brain (from posterior to anterior) with t-statistical map overlay; demonstrating effect of [A-D] injection group (Ms-PFF and Hu-PFF) compared to PBS-injected M83 mice within each sex; and [I-J] Ms-PFF compared to Hu-PFF within male and female M83 mice. Colour map describes the direction of the t-statistics; cooler colours denoting negative slopes; most commonly corresponding to volume decline and warmer colours denoting positive slopes, corresponding to volume increases over time for the group of interest indicated; t-values bounded between FDR 1 and 5% for maps (A-D) and between FDR 5 and 20% for map (J). [I] The colour map spans t-values from the max t-value to the t-critical value (significant t-value at p=0.05, prior to FDR multiple comparisons correction), increasing in opacity, the higher the t-value. On top of the colour map, contours, either black or grey, denote the voxels that meet an FDR threshold of 5% and 20% respectively. [E-H; K-N] Plot of percent relative volume change over the four time points (-7, 30, 90 and 120 days post-injection). Shading represents ±1 standard error of the mean. We chose to highlight voxels within regions where group differences over time were observed across comparisons; peak voxel in [E] periaqueductal gray (PAG), [F] right thalamus (Th), [G, M] the injection site (right caudoputamen (CP)), [H,N] right primary motor area (MC), [H,V] right hippocampus (hipp), [K] right midbrain reticular nucleus (MRN), [L] left CP. Purple line for PBS-injected mice, orange line for Hu-PFF-injected mice, green line for Ms-PFF-injected mice, solid line and triangle points for WT and dashed line and circular points for M83 hemizygous mice.

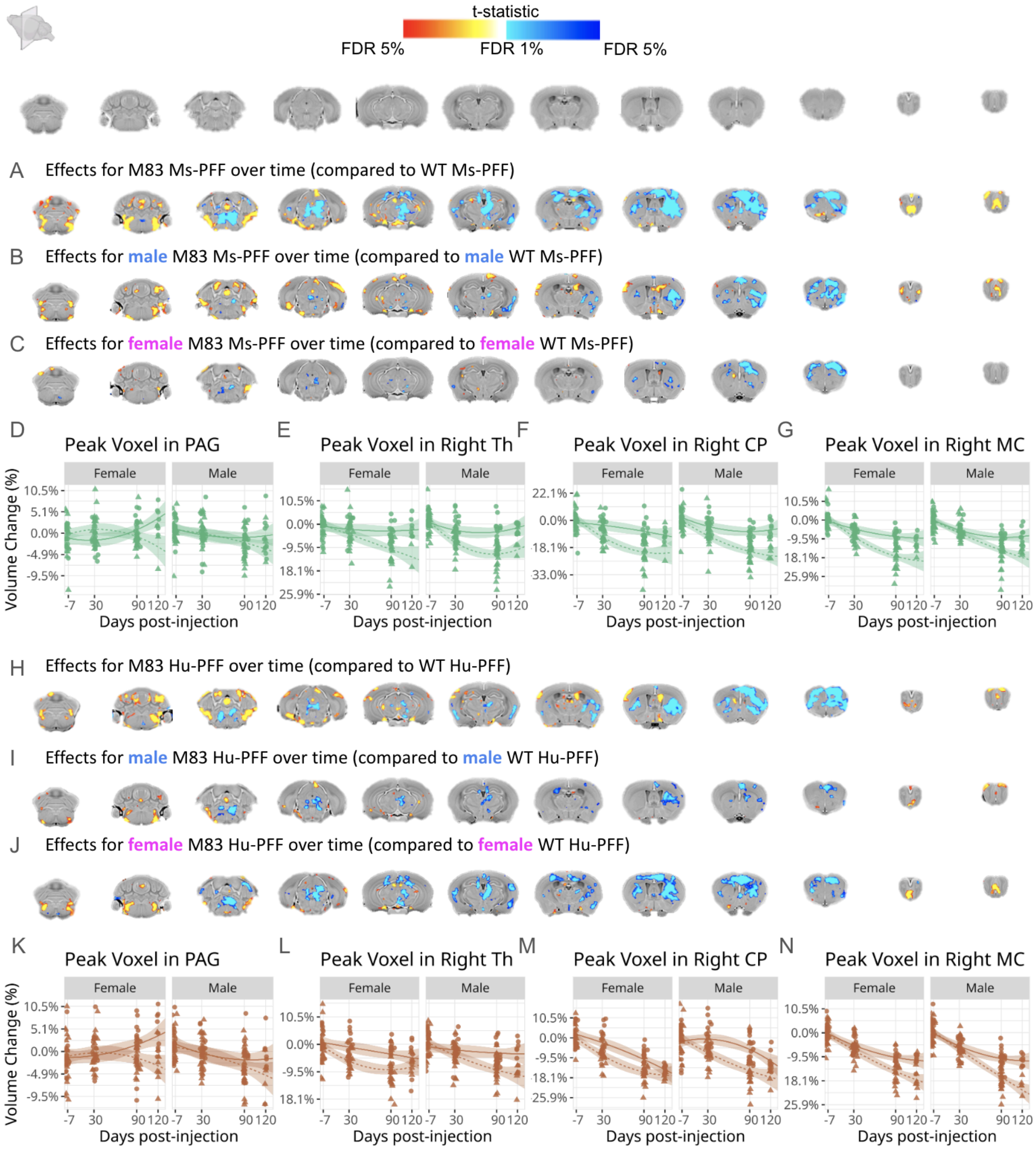


**Supplementary Figure 6. Injection group-focused analysis (with sex-based analysis) of genotype differences in voxel-wise volumetric trajectories.** [A-D] Coronal slices of the mouse brain (from posterior to anterior) with t-statistical map overlay; demonstrating effect of genotype (M83 vs WT) for each PFF injection group. The statistical differences are being shown for the [A-C] M83 Ms-PFF-injected and [H-J] M83 Hu-PFF-injected over time (compared to WT injected mice) as well as [B-C; I-J] within each sex. Colour map describes the direction of the t-statistics; cooler colours denoting negative slopes; most commonly corresponding to volume decline and warmer colours denoting positive slopes, corresponding to volume increases over time for the group of interest indicated; t-values bounded between FDR 1 and 5%. [D-G; K-N] Plot of percent relative volume change over the four time points (-7, 30, 90 and 120 days post-injection). Shading represents ±1 standard error of the mean. We chose to highlight voxels within regions where group differences over time were observed across comparisons; peak voxel in [D,K] the right side of the pons, [E,L] right thalamus (Th), [F,M] the injection site (right caudoputamen (CP)), [G,N] right primary motor area (MC). Purple line for PBS-injected mice, orange line for Hu-PFF-injected mice, green line for Ms-PFF-injected mice, solid line and triangle points for WT and dashed line and circular points for M83 hemizygous mice.

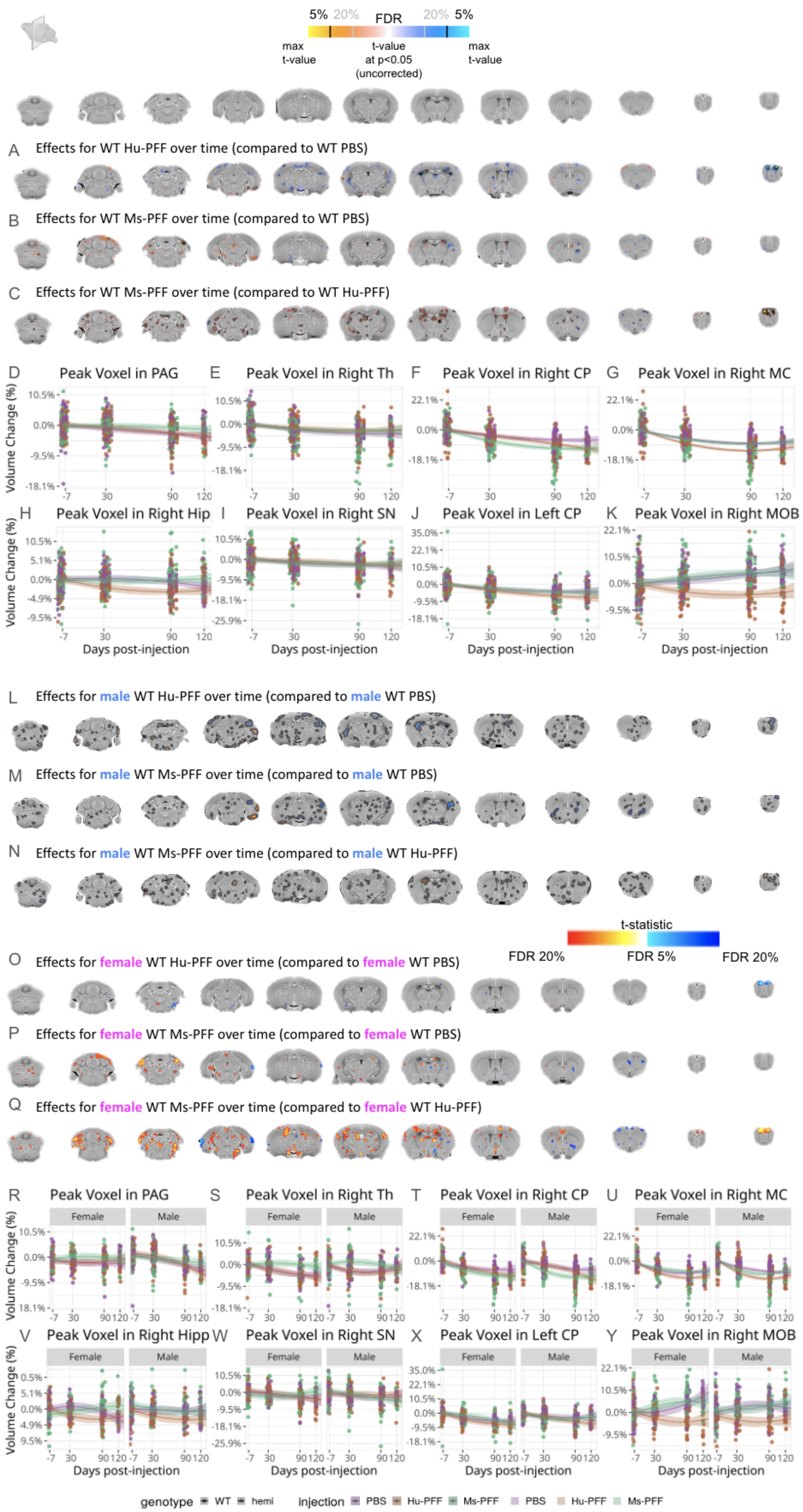
5% 20% FDR 20% 5%
max t-value
t-value at p<0.05 (uncorrected)
max t-value
A Effects for WT Hu-PFF over time (compared to WT PBS)
B Effects for WT Ms-PFF over time (compared to WT PBS)
C Effects for WT Ms-PFF over time (compared to WT Hu-PFF)
D Peak Voxel in PAG
E Peak Voxel in Right Th
F Peak Voxel in Right CP
G Peak Voxel in Right MC
H Peak Voxel in Right Hip
I Peak Voxel in Right SN
J Peak Voxel in Left CP
K Peak Voxel in Right MOB
Volume Change (%)
Days post-injection
L Effects for male WT Hu-PFF over time (compared to male WT PBS)
M Effects for male WT Ms-PFF over time (compared to male WT PBS)
N Effects for male WT Ms-PFF over time (compared to male WT Hu-PFF)
t-statistic
FDR 20% FDR 5% FDR 20%
O Effects for female WT Hu-PFF over time (compared to female WT PBS)
P Effects for female WT Ms-PFF over time (compared to female WT PBS)
Q Effects for female WT Ms-PFF over time (compared to female WT Hu-PFF)
R Peak Voxel in PAG
S Peak Voxel in Right Th
T Peak Voxel in Right CP
U Peak Voxel in Right MC
V Peak Voxel in Right Hipp
W Peak Voxel in Right SN
X Peak Voxel in Left CP
Y Peak Voxel in Right MOB
Female
Male
genotype WT hemi
injection PBS Hu-PFF Ms-PFF PBS Hu-PFF Ms-PFF

**Supplementary Figure 7. Genotype (WT) group-focused analysis (with sex-based analysis) of injection group differences in voxel-wise volumetric trajectories.** [A-C; L-N; O-Q] Coronal slices of the mouse brain (from posterior to anterior) with t-statistical map overlay; demonstrating effect of injection group (Hu-PFF compared to PBS; Ms-PFF compared to PBS; Ms-PFF compared to Hu-PFF) for WT mice. The statistical differences are being shown for the three injection group comparisons for [A-C] all subjects; [L-N] only male; and [O-Q] only female mice. Colour map describes the direction of the t-statistics; cooler colours denoting negative slopes; most commonly corresponding to volume decline and warmer colours denoting positive slopes, corresponding to volume increases over time for the group of interest indicated. Here significance was quite lower compared to the within M83 analyses (Supplementary figure 6). [A-C; L-N] The colour map spans t-values from the max t-value to the t-critical value (significant t-value at $p=0.05$, prior to FDR multiple comparisons correction), increasing in opacity, the higher the t-value. On top of the colour map, contours, either black or grey, denote the voxels that meet an FDR threshold of 5% and 20% respectively. [O-Q] The colour map spans t-values between FDR thresholds of 5% and 20%. [D-K; R-Y] Plot of percent relative volume change over the four time points (-7, 30, 90 and 120 days post-injection). Shading represents ±1 standard error of the mean. We chose to highlight voxels within regions where group differences over time were observed across comparisons; peak voxel in [D,R] periaqueductal gray (PAG), [E,S] right thalamus (Th), [F,T] the injection site (right caudoputamen (CP)), [G,U] right primary motor area (MC), [H,V] right hippocampus (hipp), [I,W] right substantia nigra (SN), [J,X] left CP and [K,Y] right main olfactory bulb (MOB). Purple line for PBS-injected mice, orange line for Hu-PFF-injected mice, green line for Ms-PFF-injected mice, solid line and triangle points for WT and dashed line and circular points for M83 hemizygous mice.

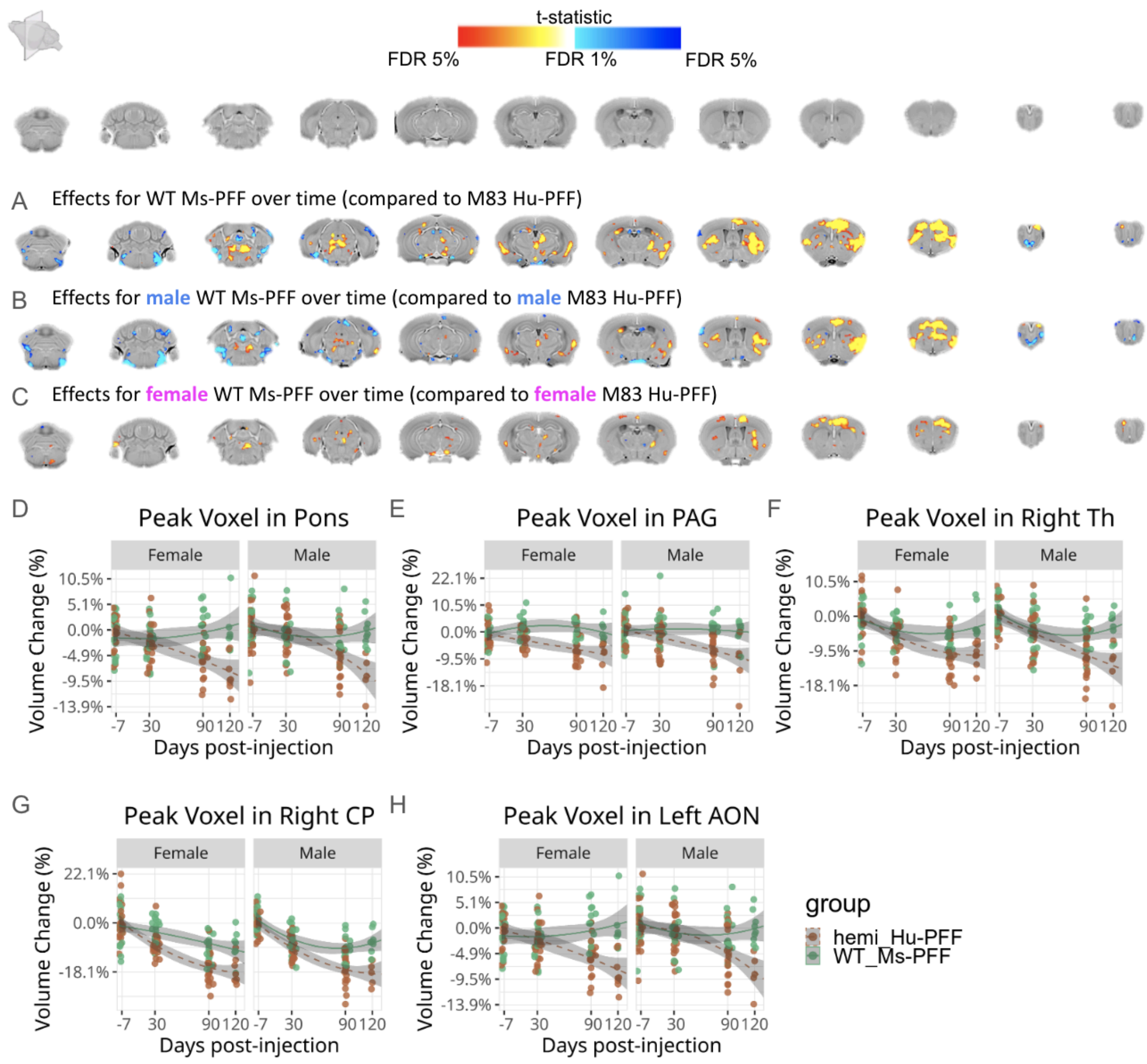


**Supplementary Figure 8. Voxel-wise differences in volume between two commonly used mouse models of synucleinopathy.** [A-C] Coronal slices of the mouse brain (from posterior to anterior) with t-statistical map overlay; demonstrating group differences between M83 Hu-PFF-injected mice and WT Ms-PFF-injected mice, as well as the group comparison within each sex. Colour map describes the direction of the t-statistics for the WT Ms-PFF-injected mice; cooler colours denoting negative slopes; most commonly corresponding to volume decline and warmer colours denoting positive slopes, corresponding to volume increases over time compared to M83 Hu-PFF-injected mice; t-values bounded between FDR 1 and 5%. [D-H] Plot of percent relative volume change over the four time points (-7, 30, 90 and 120 days post-injection). Shading represents ±1 standard error of the mean. Peak voxels displayed for the [D] right side of the pons, [E] periaqueductal gray (PAG); [F] right thalamus (Th), [G] injection site (right caudoputamen (CP)), [H] left anterior olfactory nucleus (AON). Orange line for Hu-PFF-injected mice, green line for Ms-PFF-injected mice, solid line and triangle points for WT and dashed line and circular points for M83 hemizygous mice.

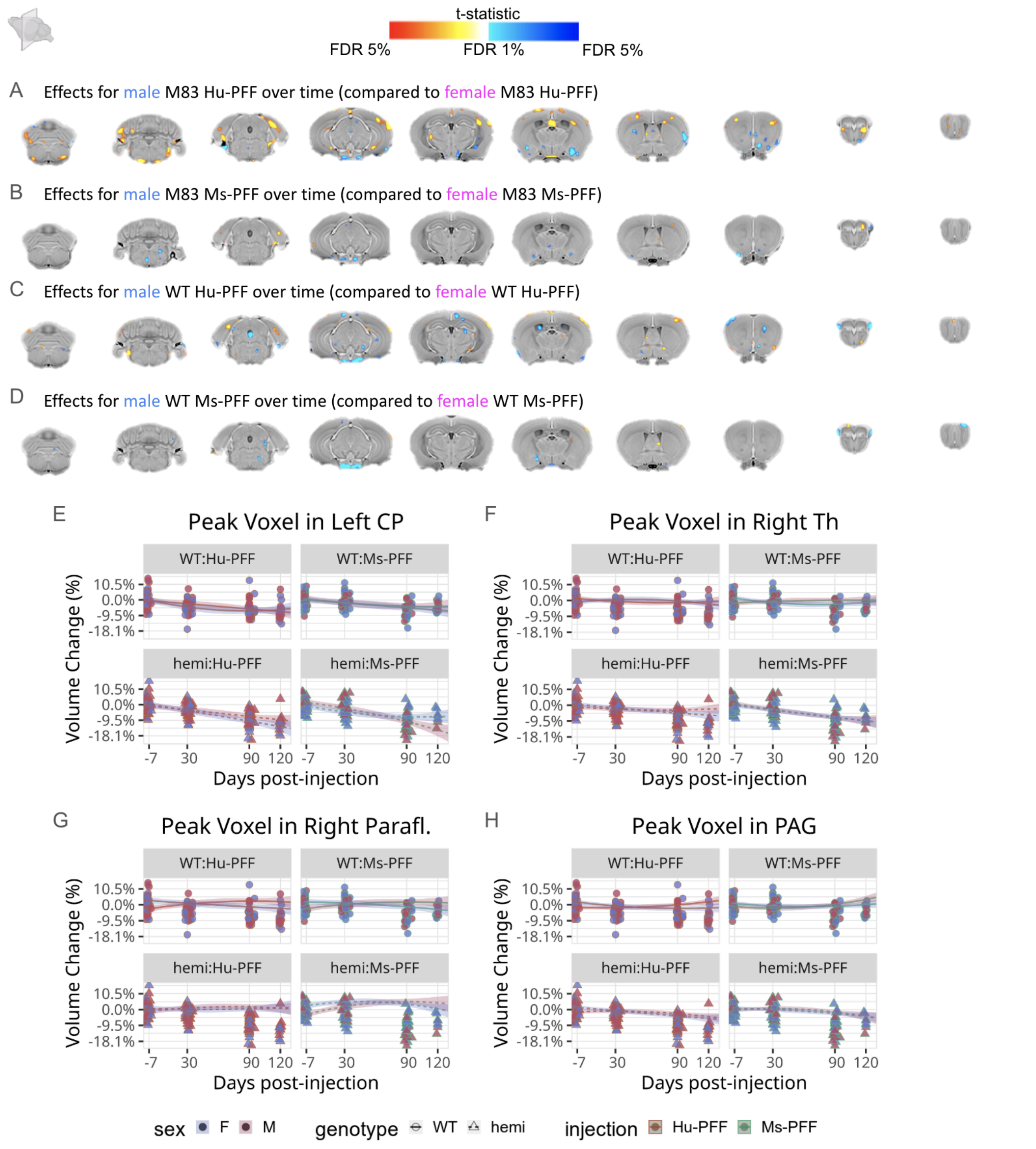


**Supplementary Figure 9. Sex differences within each PFF-injected mouse group in terms of voxel-wise volumetric trajectories.** [A-D] Coronal slices of the mouse brain (from posterior to anterior) with t-statistical map overlay; demonstrating the effects of [A] male M83 Hu-PFF compared to female M83 Hu-PFF mice over time; [B] male M83 Ms-PFF compared to female M83 Ms-PFF mice over time; [C] male WT Hu-PFF compared to female WT Hu-PFF mice over time; [B] male WT Ms-PFF compared to female WT Ms-PFF mice over time. Colour map describes the direction of the t-statistics; cooler colours denoting negative values; most commonly corresponding to volume decline and warmer colours denoting positive values, corresponding to volume increases over time for the group of interest indicated. [E-H] Plot of percent relative volume change over the four time points (-7, 30, 90 and 120 days post-injection). Shading represents ±1 standard error of the mean. Peak voxels displayed for the [E] left caudoputamen (CP); [F] right thalamus (Th); [G] right paraflocculus of the cerebellum (Paraflo.); [H] periaqueductal gray (PAG). Orange line for Hu-PFF-injected mice, green line for Ms-PFF-injected mice, red points and line shading for female and blue for males, solid line and triangle points for WT and dashed line and circular points for M83 hemizygous mice.

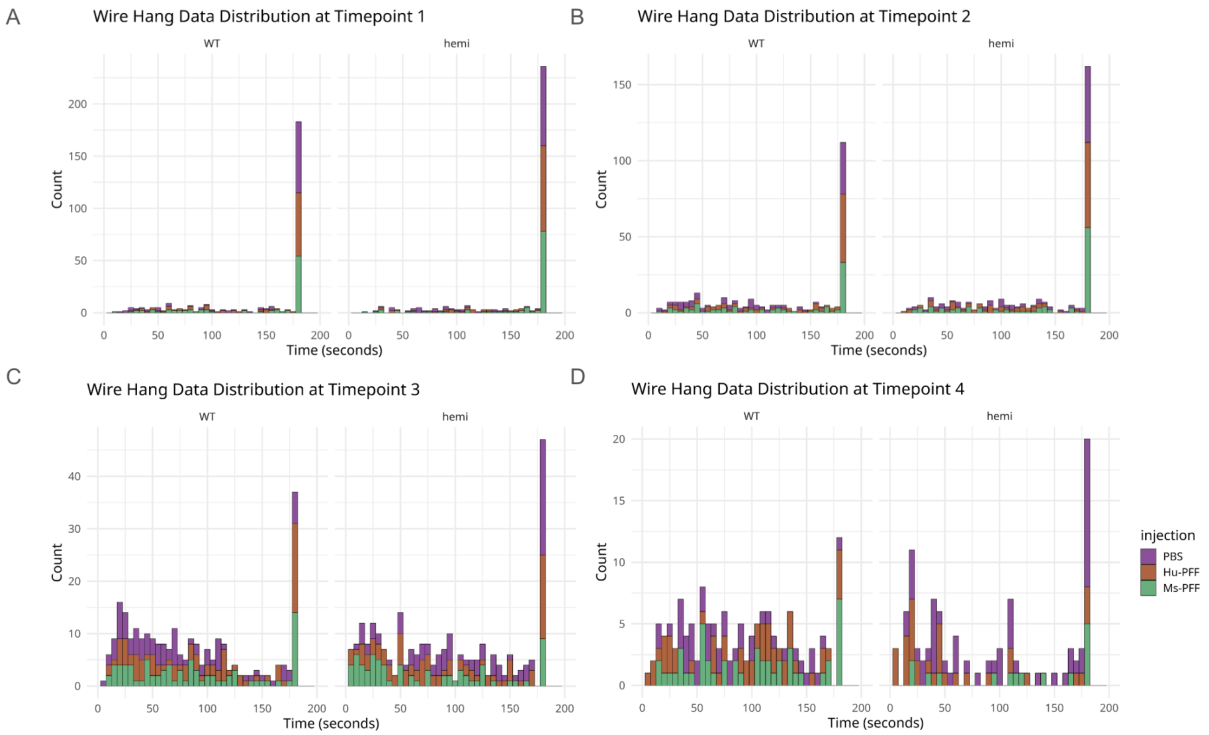


E **Proportion of Successes over time**

| | | -7 dpi | 30 dpi | 90 dpi | 120 dpi |
|---|---|---|---|---|---|
| **WT** | **PBS** | 0.6667 | 0.3474 | 0.0667 | 0.0000 |
| | **Hu-PFF** | 0.6100 | 0.4592 | 0.1954 | 0.0784 |
| | **Ms-PFF** | 0.5684 | 0.3667 | 0.1605 | 0.1458 |
| **hemi** | **PBS** | 0.6757 | 0.4804 | 0.2366 | 0.2037 |
| | **Hu-PFF** | 0.7105 | 0.5091 | 0.1613 | 0.0769 |
| | **Ms-PFF** | 0.6964 | 0.5185 | 0.1154 | 0.2778 |

**Supplementary Figure 10. Data distribution for wire hang test at each timepoint highlighting the decreasing number of mice reaching the ceiling.** [A-D] Histograms highlights the spread of the data such that there is a ceiling effect in the wire hang test, where many subjects reached the maximum allotted time for wire hang and accordingly successfully completed the test, as any latency under the 180 seconds is deemed a failed attempt. Using the Cox Proportional Hazard model, this framework provides a more accurate and robust way to analyze and interpret this type of data. This method captures both the time taken and the success/failure rates, offering a nuanced picture of performance across groups. Notably, the number of mice that successfully pass the task decreases across the time points. PBS-injected mice are in purple, Hu-PFF mice are in orange, and Ms-PFF mice are in green. Histogram binning was performed at 5. [E] Proportion of success trials at each timepoint for each mouse sorted by genotype and injection group.

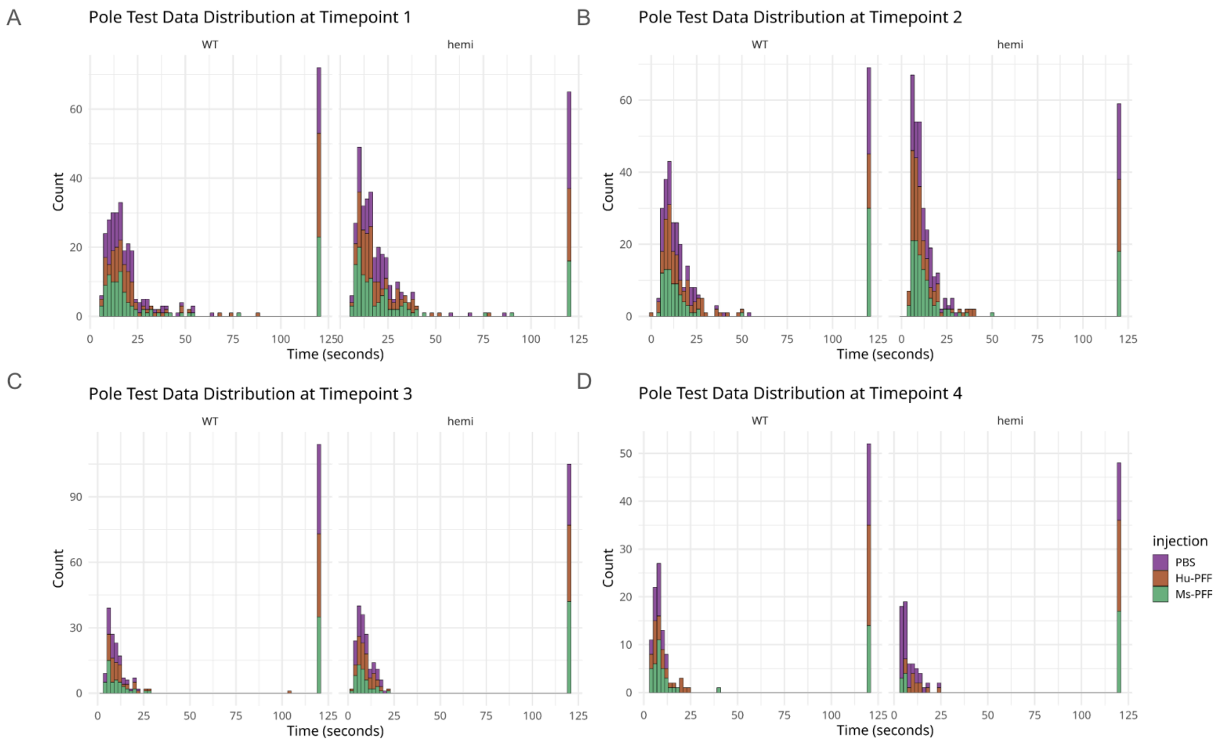


E

**Proportion of Successes over time**

| | | -7 dpi | 30 dpi | 90 dpi | 120 dpi |
|---|---|---|---|---|---|
| **WT** | **PBS** | 0.8288 | 0.7714 | 0.5444 | 0.6222 |
| | **Hu-PFF** | 0.7297 | 0.8611 | 0.5632 | 0.5882 |
| | **Ms-PFF** | 0.7870 | 0.7143 | 0.5679 | 0.7083 |
| **hemi** | **PBS** | 0.7724 | 0.8158 | 0.6989 | 0.7778 |
| | **Hu-PFF** | 0.8372 | 0.8413 | 0.6354 | 0.4242 |
| | **Ms-PFF** | 0.8760 | 0.8537 | 0.5333 | 0.2917 |

**Supplementary Figure 11. Data distribution for pole test highlights floor effect.** This histogram highlights the spread of the data such that there is a floor effect in the pole test, where many subjects failed to perform the task and consequently were allotted the maximum time (120 seconds). Using the Cox Proportional Hazard model, this framework provides a more accurate and robust way to analyze and interpret this type of data. This method captures both the time taken and the success/failure rates, offering a nuanced picture of performance across groups. Notably, the number of mice that successfully pass the task decreases across the time points . PBS-injected mice are in purple, Hu-PFF mice are in orange, and Ms-PFF mice are in green. Histogram binning was performed at 5. [E] Proportion of success trials at each timepoint for each mouse sorted by genotype and injection group.
